# Synaptic Connectivity of Sensorimotor Circuits for Vocal Imitation in the Songbird

**DOI:** 10.1101/2022.11.08.515692

**Authors:** Massimo Trusel, Ziran Zhao, Danyal H. Alam, Ethan S. Marks, Maaya Z. Ikeda, Todd F. Roberts

## Abstract

Sensorimotor computations for learning and behavior rely on precise patterns of synaptic connectivity. Yet, we typically lack the synaptic wiring diagrams for long-range connections between sensory and motor circuits in the brain. Here we provide the synaptic wiring diagram for sensorimotor circuits involved in learning and production of male zebra finch song, a natural and ethologically relevant behavior. We examined the functional synaptic connectivity from the 4 main sensory afferent pathways onto the 3 known classes of projection neurons of the song premotor cortical region HVC. Recordings from hundreds of identified projection neurons reveal rules for monosynaptic connectivity and the existence of polysynaptic ensembles of excitatory and inhibitory neuronal populations in HVC. Circuit tracing further identifies novel connections between HVC’s presynaptic partners. Our results indicate a modular organization of ensemble-like networks for integrating long-range input with local circuits, providing important context for information flow and computations for learned vocal behavior.

## Introduction

Patterns of synaptic connectivity anchor how activity flows within sensory and motor networks in the brain. Consequently, these synaptic connections delimit and help define how sensory and premotor circuits interact. Mapping synaptic pathways between these circuits is therefore fundamental to understanding the neural computations involved in learning and producing behaviors. Great strides have been made in developing technologies for large-scale recording of neuronal activity across brain regions^1-3^ and in building functional connectomes of local (< 1 mm) synaptic circuits^4^. Less progress has been made at intermediate levels, which aim to build wiring diagrams for long-range synaptic connections in the brain^5^. Moreover, while most evidence for long range synaptic connectivity has been provided piecemeal, we now have the opportunity to systematically apply circuit dissection methods to other animal models in which we still know relatively little about patterns of long-range synaptic connectivity.

We were inspired to understand the wiring diagram for sensorimotor circuits critical for birdsong, a complex and volitionally produced skilled behavior that, like speech and language, is dependent on forebrain circuits for its fluent production^6,7^. The dedicated neural circuits associated with song provide a powerful model in which to study how synaptically linked sensory and motor networks of neurons control a complex behavior. The courtship song of male zebra finches is one of the better studied naturally learned behaviors^8-20^. Zebra finch song is controlled by a relatively discrete set of interconnected forebrain regions located in the dorsal ventricular ridge (DVR)^21-30^. The DVR is the avian analogue of the mammalian neocortex, and like the neocortex it contains parcellated regions that (i) receive sensory information from the thalamus, (ii) regions that process information connected by mostly ipsilaterally projecting intratelencephalic projection neurons (PNs), and (iii) output motor circuits projecting back to the thalamus and to motor regions throughout the brainstem^31,32^. The vocal premotor nucleus HVC (proper name) is a central hub in the DVR song network. HVC is necessary for juvenile song learning and adult song production. It also serves as the primary synaptic interface between sensory pathways and the premotor circuits involved in learning and controlling singing behavior^21,27,28,33-36^.

Although anatomical evidence delineated the main input and output pathways of HVC decades ago^21,27,37-40^, a cell-type specific synaptic wiring diagram of this core circuitry has remained out of reach. HVC receives intratelencephalic input from at least 4 sensory and sensorimotor regions in the DVR (NIf, Av, mMAN and RA) and one thalamic brain region, Uva (**Fig. 1A, B**; see Fig.1 legend for anatomical descriptions). Several lines of evidence support the idea that song learning, and vocal motor control involves some of these pathways projecting into HVC^27,34,36,40-46^. In addition, HVC has 3 distinct classes of intratelencephalic projecting neurons that function as the output pathways important for song learning and song motor control ^27^. HVC’s projection onto RA through HVC_RA_ neurons forms the descending song motor pathway necessary for song production. HVC’s projection onto Area X through HVC_X_ neurons forms a DVR-striatal circuit that is important for song plasticity. HVC’s projection onto Avalanche through HVC_Av_ neurons forms the song circuit’s projection back to the auditory DVR, a pathway that is important for evaluating motor performance during song learning^27^.

**Fig. 1:**
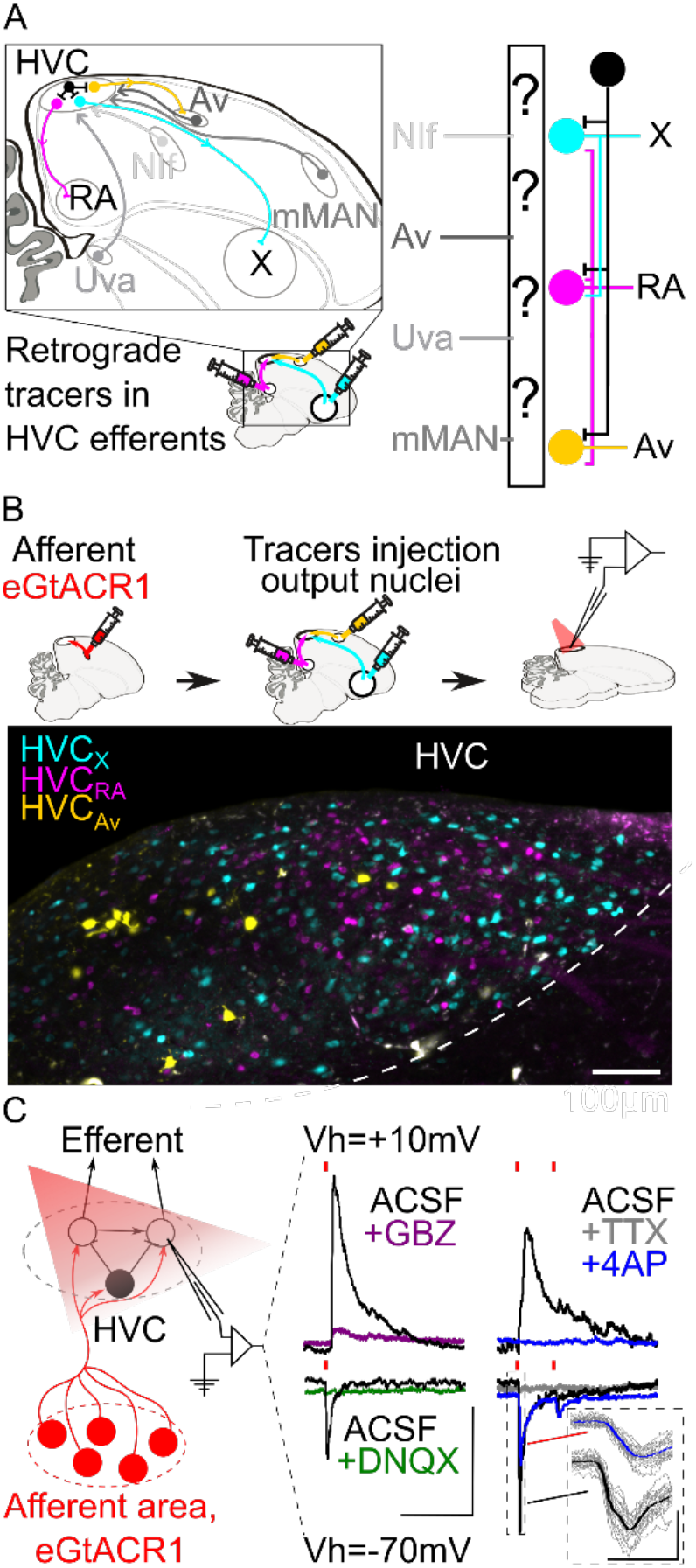
Strategy for opsin assisted HVC circuit mapping. **A)** (left) Parasagittal schematic of the zebra finch brain illustrating HVC afferents (grey) from nucleus interface of the nidopallium (NIf), nucleus avalanche (Av), medial magnocellular nucleus of the anterior nidopallium (mMAN) and nucleus Uvaeformis (Uva), and the 3 classes of HVC-PNs: HVC_X_ (projecting to Area X, cyan), HVC_RA_ (projecting to RA, magenta) and HVC_Av_ (projecting to Av, yellow). (right) Schematic illustrating the known synaptic connectivity of HVC’s input and output pathways (interneurons schematized in black). **B**) (top) Schematic of the workflow including opsin expression in afferent areas, retrograde tracer injection in afferent areas and whole cell patch-clamp recording of light-evoked currents in acute brain slices. (bottom) Sample image of retrogradely labeled HVC-PN classes in a brain slice used for patch-clamp recordings (scalebar 100µm). **C**) Schematic of whole-cell recording of light evoked synaptic currents in HVC-PNs receiving monosynaptic inputs from one of the afferent areas expressing eGTACR1 (red), as well as polysynaptic inputs from local HVC-PNs (white circle) and interneurons (black circle). Recordings are performed at holding potential (Vh)= +10mV and - 70mV. Sample traces report the effect of bath application of DNQX (green) and gabazine (GBZ, purple), which suppress oEPSC and oIPSC, respectively. Glutamatergic monosynaptic currents are pharmacologically isolated by bath application of TTX+4AP: sample traces (average of 20 sweeps) portray a typical case of a cell displaying oEPSC with a monosynaptic component (inset shows 20 oEPSCs sweeps in grey, averaged in the thick traces in black and blue), and polysynaptic oIPSC, as 4AP (blue) application results in a partial restoration of oEPSC, but not of oIPSC (red lines represent light stimuli, 1ms; scalebar: 100ms, 100pA).

Mapping the synaptic connectivity between HVC’s input and output pathways has been hindered by the lack of tools for robustly manipulating these circuits. We have overcome this issue by employing better performing optogenetic channels packaged in an adeno associated virus (AAV) that we optimized for expression throughout various portions of the song circuitry. Specifically, we made use of the axonal expression of the opsin eGtACR1^47,48^. GtACRs are blue light-driven Cl^-^ channels that, while hyperpolarizing at the soma and dendrites, are depolarizing at axon terminals due to the shifted internal Cl^-^ concentration in axonal compartments^49-51^. Here we use viral expression of eGtACR1 to achieve independent control of synaptic release from NIf, Uva, mMAN, and Av. We paired this with whole-cell patch-clamp recordings from visually targeted HVC projection neurons (HVC-PNs; HVC_X_, HVC_RA_, HVC_Av_), identified using retrograde tracer injections into each efferent region (**Fig. 1C**). This permitted the first large-scale cell-type specific synaptic connectivity mapping, using gold-standard electrophysiological methods, in songbirds. In ex-vivo brain slices from adult male zebra finches, we mapped the properties of synaptic transmission at the 12 afferent-HVC-PN combinations. Via recordings from hundreds of HVC PNs, we provide a comprehensive polysynaptic and monosynaptic mapping of the connectivity between these afferent sensory pathways and the output pathways of HVC. Through this research, we provide the first long-range cellular resolution synaptic connectivity map in the avian DVR, a fundamental step in the effort to identify common themes and differences in synaptic connectivity between the DVR and mammalian neocortical circuits.

## Results

To conduct opsin-assisted circuit mapping in zebra finches, we expressed eGtACR1 in NIf, Uva, mMAN, or Av using AAVs (Fig. **1C**; a cocktail of AAV2.9-Cbh-FLP and AAV2.9-Cbh-fDIO-eGtACR1 injected in HVC afferent areas). 4-6 weeks later we injected retrograde tracers in HVC projection targets (RA, Area X, and Av) to label HVC-PN classes in different fluorescent channels. After 2-7 days we obtained acute brain slices and performed whole-cell patch-clamp recordings from visually identified HVC-PNs to examine optically evoked post-synaptic currents (oPSCs). We confirmed that oPSCs measured while holding the membrane voltage at -70mV were excitatory (oEPSCs) and were mediated by AMPA receptors (current suppressed by bath application of DNQX; Fig. **1C**). Holding the membrane potential to +10mV allowed us to measure inhibitory currents (oIPSCs) mediated by GABAa receptors (current suppressed by bath application of gabazine; Fig. **1C**).

For each pathway we report the likelihood of observing oEPSCs and oIPSCs, the amplitude of oEPSCs and oIPSCs, and the excitatory-inhibitory ratio (oEPSC/oIPSC). Although we calculated the paired-pulse ratio, the slow kinetics of eGtACR1 make interpretation of these results difficult, therefore it is not being reported. We also report whether observed synaptic currents have a monosynaptic component, by applying the voltage-gated Na^+^ channel blocker tetrodotoxin (TTX) followed by the K^+^ channel blocker 4-aminopiridyne (4AP)^5,52^ (Fig. **1C**). TTX blocks action potentials and the addition of 4AP blocks the rapid K+ dependent repolarization of the axon. This allows local opsin-driven depolarizations to reach threshold for calcium-dependent vesicle docking and release, thus revealing optically evoked monosynaptic currents^52^. We consistently observed that, while oEPSCs registered in ACSF displayed multiple peaks consistent with a polysynaptic source, currents suppressed by TTX that returned following bath application of 4AP had a monotonic rising slope and a single peak, consistent with a monosynaptic origin of the current (Fig. **1C**).

### Uva monosynaptically innervates HVC_RA_ neurons

Nucleus Uvaeformis (Uva) is a small polysensory nucleus in the caudal thalamus that is reported to be necessary for song production. Uva is a recipient of somatosensory information from the trigeminal system, visual input from the optic tectum, auditory information from the lateral lemniscus, cholinergic input from the medial habenula, and information about respiratory timing from the ventral respiratory column ^53-58^. Its inputs to HVC are thought to be pivotal in song motor control ^29,42,43^. However, how Uva synaptically influences HVC activity is still poorly understood. We found that Uva neurons are robustly transduced by our AAV expressing eGtACR1 (**Fig. 2A**). Thalamic regions around Uva do not project to HVC, allowing us to selectively map Uva’s input to the 3 classes of HVC-PNs.

**Fig. 2:**
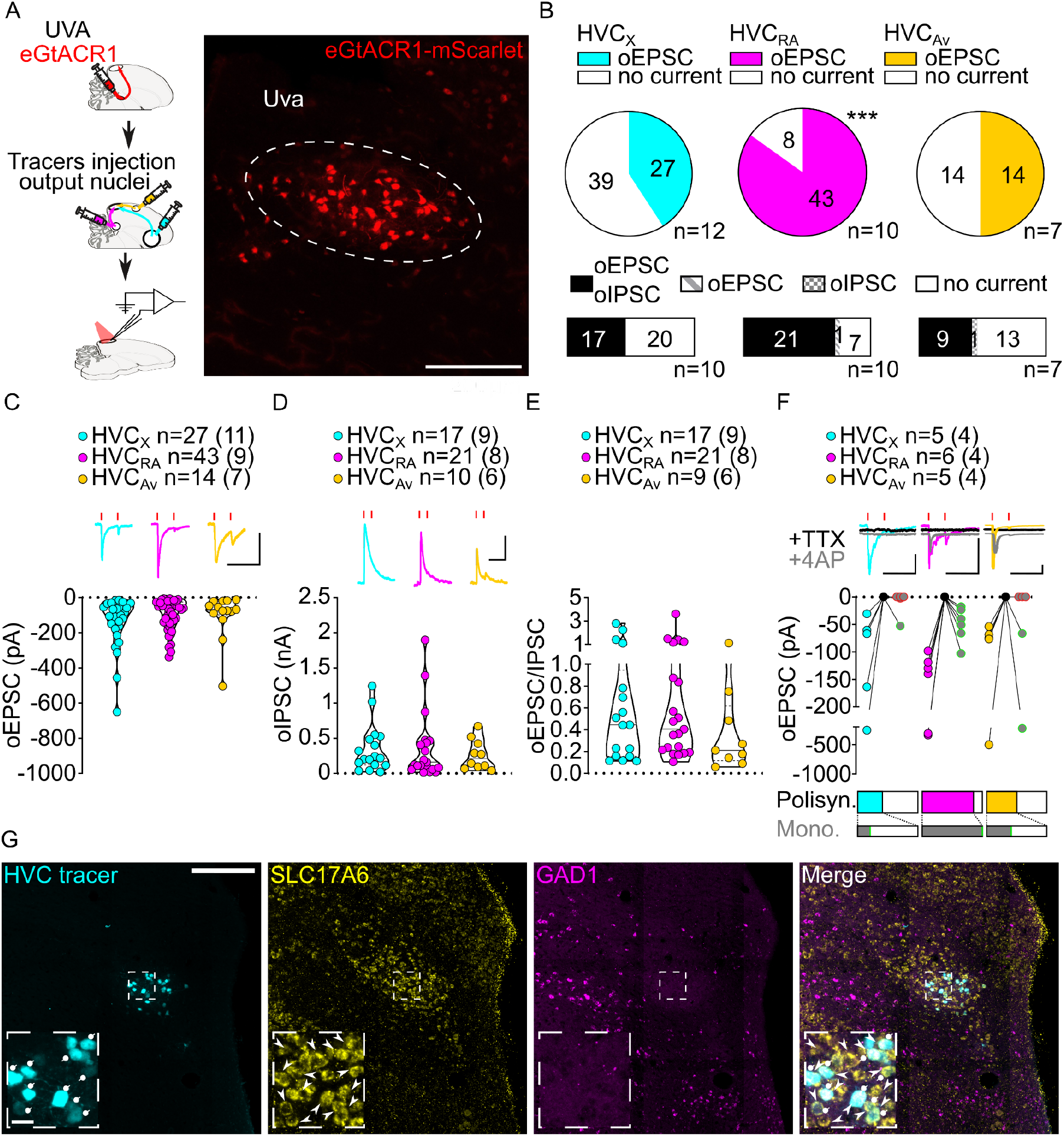
Uva afferents reliably elicit polysynaptic oIPSCs and oEPSCs in all three HVC-PN classes and have monosynaptic inputs onto HVC_RA_ neurons. **A**) (left) Schematic of the experimental timeline, illustrating injection of AAV-eGtACR1 in Uva, followed by retrograde tracer injections in HVC efferent areas and whole-cell patch-clamp recording in acute brain slices; (right) sample image of eGtACR1-mScarlet expression in Uva (scalebar 200µm). **B**) (top) pie charts representing the likelihood of observing oEPSCs in HVC_X_ (cyan), HVC_RA_ (magenta) or HVC_Av_ (yellow) (Fisher’s exact test, P<0.001). Numbers in the pie fragments represent the number of cells in which current (colored) or no current (white) was found. Numbers next to the pie charts represent the number of animals from which the data is obtained. (bottom) bar chart representing the number of cells where both oEPSCs and oIPSCs could be elicited (black), only oEPSCs but no oIPSCs (grey lines), only oIPSCs but no oEPSCs (grey checkers), or neither (white), for subsets of cells from the HVC-PN classes’ pie charts aligned above (Fisher’s exact test, P=0.0244). **C**) violin and scatter plot and sample traces reporting average measured oEPSC amplitude of each cell, by cell class (Kruskal-Wallis test H(2)=2.241, P=0.3262; n= cells (animals); red lines represent light stimuli, 1ms; scalebars, 100ms, 100pA). **D**) violin and scatter plot and sample traces of average measured oIPSC amplitude of each cell, by cell class (H(2)=0.5946, P=0.7428). **E**) violin and scatter plot of the ratio of oEPSC and oIPSC peak amplitude of each cell where both are measured and ≠0, per cell class (H(2)=0.2716, P=0.2572). **F**) (top) sample traces and plot representing the amplitude of post-synaptic currents evoked by lightly-driven release of neurotransmitter from Uva axonal terminals in HVC; oEPSCs amplitudes are reported before (HVC_X_ cyan, HVC_RA_ magenta, HVC_Av_ yellow) and after bath application of TTX (black) and 4AP (grey, green outline indicates polysynaptic oEPSC, see methods), (n= cells (animals); blue lines represent light stimuli, 1ms; scalebars, 100ms, 100pA) (bottom) bar charts representing the likelihood of observing polysynaptic oEPSCs in HVC_X_ (cyan), HVC_RA_ (magenta) or HVC_Av_ (yellow) (data from panel B), and (grey) likelihood of a subset of the corresponding oEPSCs to be monosynaptic. **G**) Sample images reporting retrogradely labeled HVC-projecting neurons in UVA (cyan, white circles) together with in-situ labeling of glutamatergic (SLC17A6, yellow, white arrowheads) and GABAergic (GAD1, magenta, gray arrowheads) markers transcripts (scalebar 200µm, inset 20µm).

In acute brain slices we observed glutamatergic oEPSCs in all 3 classes of retrogradely identified HVC-PNs (27/66 HVC_X_, 43/51 HVC_RA_, 14/28 HVC_Av_ neurons; **Fig. 2B, Suppl. Fig. 1**). Optical stimulation of Uva axons resulted in significantly lower probability of observing oEPSCs in HVC_X_ and HVC_Av_ than in HVC_RA_ neurons (**Fig. 2B, Suppl. Fig. 1**). When we held the membrane voltage to +10mV we also observed relatively delayed GABAergic currents (53.93% probability, **Suppl. Fig. 1**), consistent with disynaptic inhibition. Uva axon stimulation elicited oIPSCs mainly in cells where we observed oEPSCs (oIPSC contingent with oEPSC: HVC_X_, 100.0%, HVC_RA_ 96.6%, HVC_Av_, 95.8%; **Fig. 2B**).

Despite the markedly higher probability of eliciting currents in HVC_RA_ neurons, oEPSC and oIPSC amplitudes were comparable across all the HVC-PN classes (**Fig. 2C, D, Suppl. Fig. 1**). The excitation/inhibition ratio revealed that Uva terminal stimulation generally led to higher amplitude oIPSCs than oEPSCs (**Fig. 2E, Suppl. Fig. 1**).

We next examined if HVC-PN classes receive monosynaptic input from Uva. Recent studies have suggested different views for how Uva may influence the propagation of activity in HVC. Uva has been proposed to provide high frequency (30 – 60Hz) synchronous synaptic inputs that facilitate activity propagation across HVC_RA_ premotor neurons^45^. In another proposal, Uva selectively drives activity in only a fraction, ∼16%, of HVC_RA_ neurons which are active during syllable transitions ^29^. Consistent with the higher likelihood of polysynaptic excitation in HVC_RA_ neurons, we found that oEPSC returned after application of TTX+4AP in all HVC_RA_ neurons tested (**HVC**_**RA**_: 6/6 neurons) while HVC_X_ and HVC_Av_ were infrequently found to receive monosynaptic input from Uva (HVC_X_: 1/5 cells, HVC_Av_: 2/5 cells, **Fig. 2F**). Our synaptic circuit mapping provides the first conclusive evidence that Uva makes monosynaptic connections with HVC_RA_ neurons. Given that we observed oEPSCs in 84.3% of recorded HVC_RA_ and found monosynaptic connections from Uva onto 100% of the HVC_RA_ neurons tested, our data further suggests that Uva is unlikely to make monosynaptic connections with only a small percentage of HVC_RA_ neurons.

Previous studies have been equivocal to the excitatory, inhibitory, or neuromodulatory nature of the Uva to HVC circuitry _29,53_. We did not observe monosynaptic oIPSCs in any cell class suggesting that Uva provides only a glutamatergic input to HVC (HVC_X_: 0/3, HVC_RA_: 0/5, HVC_Av_: 0/4, data not shown). We combined retrograde labeling from HVC with in-situ hybridization labeling on brain slices. This revealed that Uva neurons projecting to HVC (Uva_HVC_) selectively expressed the glutamatergic marker SLC17A6, but not the GABAergic marker GAD1^32^ (**Fig. 2G)**. Interestingly, our labeling did not identify any GABAergic neurons within Uva, indicating that Uva_HVC_ may function as an excitatory relay of diverse polysensory input pathways. Although we cannot exclude the possibility that Uva neurons may also co-release neuropeptides, our results define the excitatory synaptic connectivity between Uva and HVC PNs.

**Suppl. Fig 1 (related to Fig. 2).**
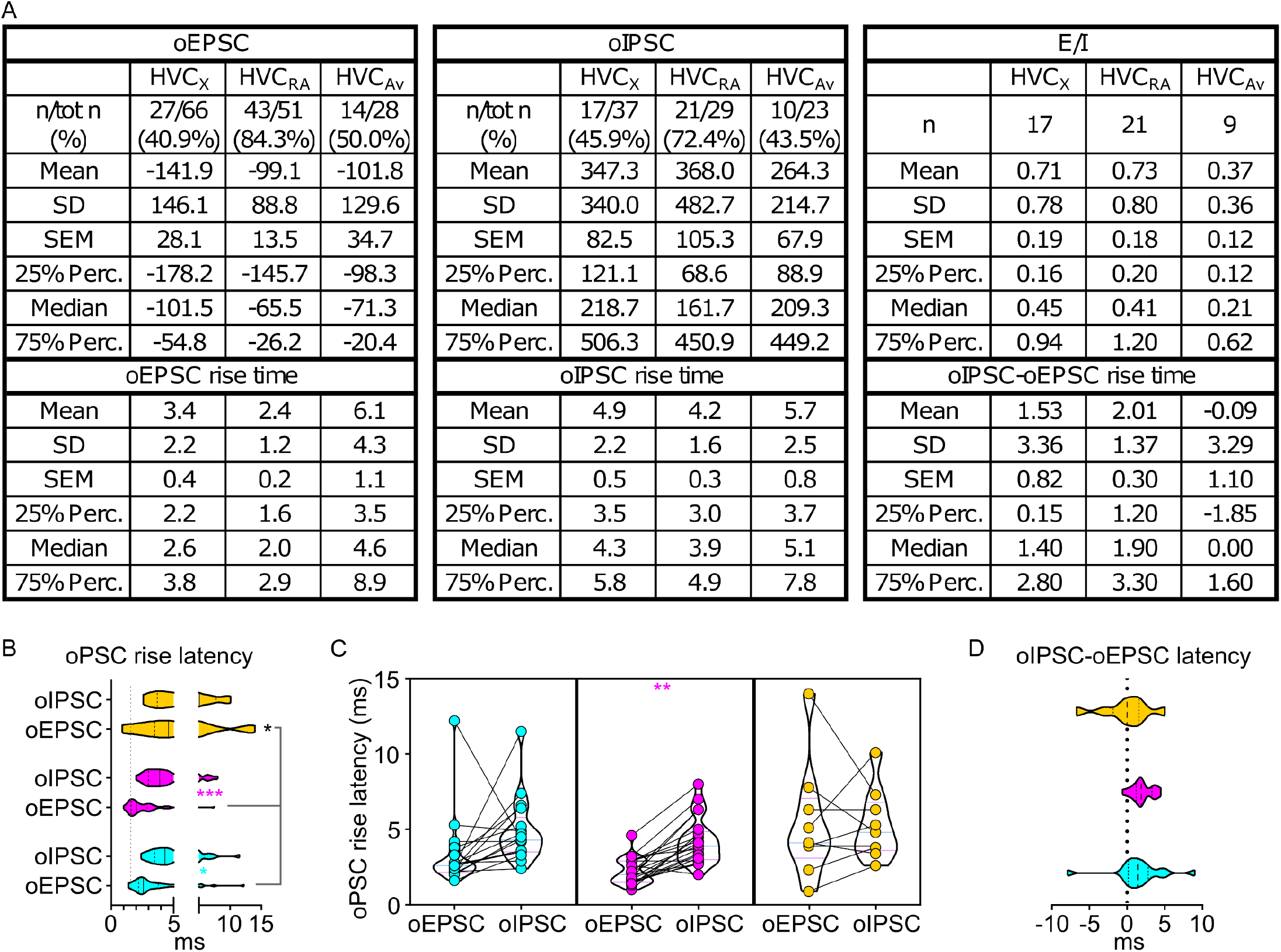
**A**) Table reporting descriptive statistics relative to oEPSCs and oIPSCs as reported in Fig. 2. **B)** violin and scatter plots representing oPSC rise latencies reported in fig. 2 (gray line marks the minimal latency (1_st_ quartile) to oPSC rise across the cell types, for ease of comparison (Mixed-effects analysis, oEPSC vs. oIPSC F(1,44)=7.143, P=0.0105 HVCX P=0.0164, HVCRA P<0.001; HVCPN F(2,82)=10.24, P<0.001, oEPSCs:HVC_X_ vs HVC_Av_ P<0.001, HVC_RA_ vs HVC_Av_ P<0.001); **C**) oPSCs rise latency for cells where both oEPSC and oIPSC were measured (2W ANOVA, oEPSC vs. oIPSC F(1,44)=7.923, HVCRA P=0.0032); **D**) relative delay of oIPSC compared to oEPSC across the cells reported in C) (Kruskal-Wallis test, H=4.587, P=0.1009)

### NIf provides excitatory monosynaptic input to all HVC-PNs

We next examined the synaptic transmission between the nucleus NIf and the 3 classes of HVC-PNs. NIf is a higher order polysensory DVR nucleus located adjacent to the primary auditory DVR (Field L2 - region that receives thalamic input from the avian homologue of the medial geniculate nucleus). NIf is recipient of afferents from Uva and multiple areas of the auditory DVR and has an essential role in relaying auditory signals to HVC^59^. The synaptic connection between NIf and HVC is obligatory in a young male zebra finch’s ability to form a memory of their father’s song^34,46^. This memory is used as birds evaluate their song performances, permitting vocal imitation of song^33^. Optogenetic manipulations of NIf inputs to HVC are also sufficient to implant song memories that guide song syllable imitation^34^. Although NIf is selectively active during singing, it is not necessary for producing song in adulthood or the song imitation process once the memory of a father’s song is acquired in juvenile birds ^34,46,60,61^.

To map synaptic connections from NIf to HVC, we expressed eGtACR1 in NIf. While the surrounding area, Field L, has been described to project to the ventral boarder of HVC^38,62,63^, we recorded from visually identified, retrogradely labeled HVC projection neurons. Area Avalanche is the only other nearby brain region that also sends projections to HVC and we carefully checked for a lack of expression in this region in all brain hemispheres used in this study (**Fig. 3A**). This provides confidence that the currents we measured are evoked by stimulating neurotransmission from NIf terminals. Light stimulation (1 ms) reliably elicited glutamatergic oEPSCs in all three classes of HVC projection neurons (78.6% probability, **Fig. 3B, Suppl. Fig. 2**). We examined oIPSCs in a subset of neurons and found strong GABAergic currents evoked by optical stimulation in all three classes of projection neurons (65.9% probability, **Fig. 3B, Suppl. Fig. 2**). oIPSCs were predominantly found in cells in which we also observed oEPSCs (oIPSC contingent with oEPSC: HVC_X_ 90.0%; HVC_RA_ 76.5%, HVC_Av_ 94.1%; **Fig. 3B**).

**Fig. 3:**
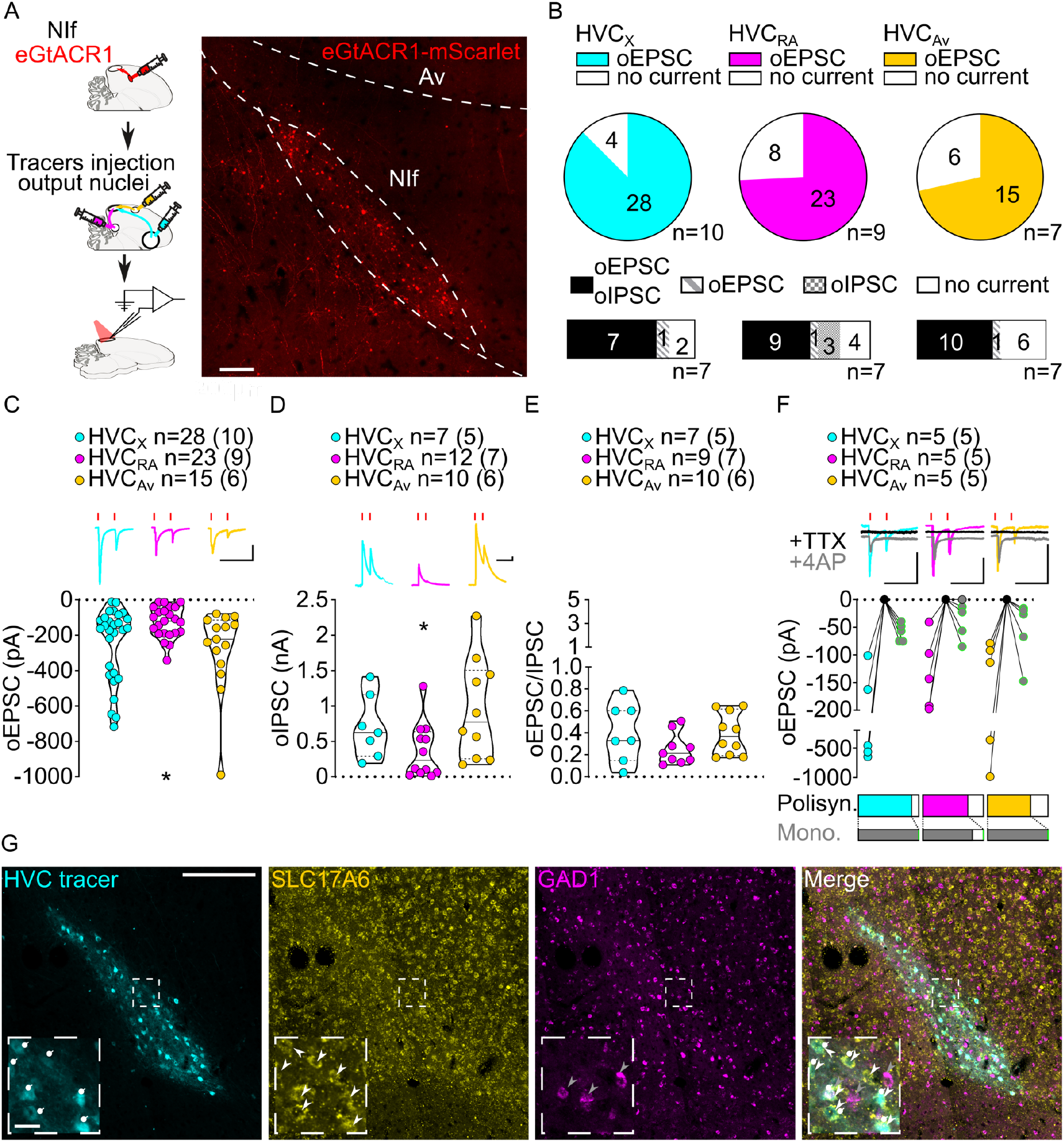
Stimulation of NIf inputs to HVC reliably elicits polysynaptic oIPSCs and oEPSCs, and monosynaptic oEPSCs, in all three HVC-PN classes. **A**) (left) Schematic of the experimental timeline, illustrating injection of AAV-eGtACR1 in NIf, followed by retrograde tracer injections in HVC efferent areas and whole-cell patch-clamp recording in acute brain slices; (right) sample image of eGtACR1-mScarlet expression in NIf (scalebar 200µm). **B**) (top) pie charts representing the likelihood of observing oEPSCs in HVC_X_ (cyan), HVC_RA_ (magenta) or HVC_Av_ (yellow) (Fisher’s exact test, P=0.2783). Numbers in the pie fragments represent the number of cells in which current (colored) or no current (white) was found. Numbers next to the pie charts represent the number of animals from which the data is obtained. (bottom) bar chart representing the number of cells where both oEPSCs and oIPSCs could be elicited (black), only oEPSCs but no oIPSCs (grey lines), only oIPSCs but no oEPSCs (grey checkers), or neither (white), for subsets of cells from the HVC-PN classes’ pie charts aligned above (Fisher’s exact test, P=0.5841). **C**) violin and scatter plot and sample traces reporting average measured oEPSC amplitude of each cell, by cell class (Kruskal-Wallis test, H(2)=6.135, P=0.0465; n= cells (animals); red lines represent light stimuli, 1ms; scalebars, 100ms, 100pA). **D**) violin and scatter plot and sample traces of average measured oIPSC amplitude of each cell, by cell class (H(2)=6.182, P=0.0455). **E**) violin and scatter plot of the ratio of oEPSC and oIPSC peak amplitude of each cell where both are measured and ≠0, per cell class (H(2)=3.305, P=0.1916). **F**) (top) sample traces and plot representing the amplitude of post-synaptic currents evoked by lightly-driven release of neurotransmitter from NIf axonal terminals in HVC; oEPSCs amplitudes are reported before (HVC_X_ cyan, HVC_RA_ magenta, HVC_Av_ yellow) and after bath application of TTX (black) and 4AP (grey, green outline indicates polysynaptic oEPSC, see methods), (n= cells (animals); blue lines represent light stimuli, 1ms; scalebars, 100ms, 100pA) (bottom) bar charts representing the likelihood of observing polysynaptic oEPSCs in HVC_X_ (cyan), HVC_RA_ (magenta) or HVC_Av_ (yellow) (data from panel B), and (grey) likelihood of a subset of the corresponding oEPSCs to be monosynaptic. **G**) Sample images reporting retrogradely labeled HVC-projecting neurons in NIf (cyan, white circles) together with in-situ labeling of glutamatergic (SLC17A6, yellow, white arrowheads) and GABAergic (GAD1, magenta, gray arrowheads) markers transcripts (scalebar 200µm, inset 20µm).

We found oEPSC and oIPSC amplitudes to be comparable between HVC_X_ and HVC_Av_ PNs (**Fig. 3C, 3D, Suppl. Fig. 2**). However, HVC_RA_ neurons had lower amplitude currents than the other HVC-PN classes (**Fig. 3C, 3D, Suppl. Fig. 2**). When we computed the amplitude of oEPSCs and oIPSCs for each cell to calculate the excitation/inhibition, we observed that optogenetic stimulation of NIf axonal terminals evoked relatively higher amplitude oIPSCs than oEPSCs in all HVC-PN classes (**Fig. 3E, Suppl. Fig. 2**).

We next examined monosynaptic connectivity from NIf to each HVC-PN. We found that application of TTX followed by 4AP reliably returned oEPSCs in all three HVC projection subtypes (monosynaptic oEPSCs: HVC_X_: 5/5, HVC_RA_: 4/5, HVC_Av_: 5/5, **Fig. 3F**). This widespread monosynaptic connectivity is in stark contrast to thalamic excitatory inputs from Uva that preferentially target HVC_RA_ neurons. Similar to Uva projections however, we did not observe monosynaptic oIPSCs (HVC_X_: 0/5, HVC_RA_: 0/2, HVC_Av_: 0/3, data not shown), suggestive of a glutamatergic projection from NIf to HVC. We combined retrograde labeling from HVC with in-situ hybridization labeling on brain slices. NIf neurons projecting to HVC (NIf_HVC_) selectively expressed the glutamatergic marker SLC17A6, but not the GABAergic marker GAD1^32^ (**Fig. 3G)**. This finding is consistent with previous studies indicating that NIf projections result in disynaptic inhibition onto HVC projection neurons^64^.

Together, these results indicate that intratelencephalic DVR circuits are excitatory and that NIf provides excitatory glutamatergic monosynaptic projections onto all 3 known classes of HVC-PNs. Given that monosynaptic connectivity across regions of the DVR have not previously been examined, this suggests the possibility that, in contrast to the thalamo-DVR pathways, projections between regions of the DVR might not have cell type selectivity.

**Suppl. Fig 2 (related to Fig. 3).**
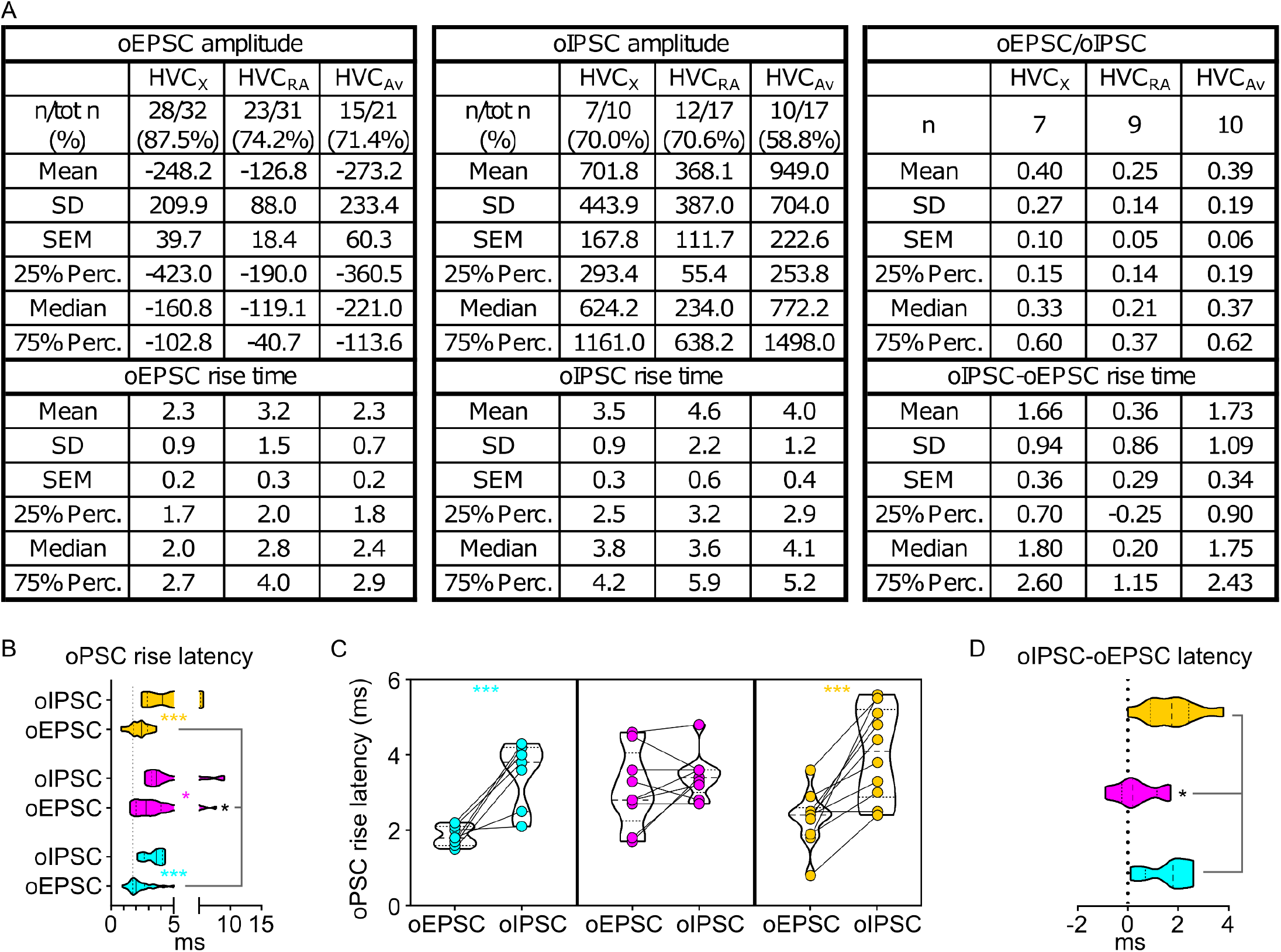
**A**) Table reporting descriptive statistics relative to oEPSCs and oIPSCs as reported in Fig. 3. **B)** violin and scatter plots representing oPSC rise latencies reported in fig. 3 (gray line marks the minimal latency (1_st_ quartile) to oPSC rise across the cell types, for ease of comparison (Mixed-effects analysis, oEPSC vs. oIPSC F(1,23)=46.55, P<0.001 HVC_X_ P<0.001, HVC_RA_ P=0.0167. HVC_Av_ P<0.001; HVC_PN_ F(2,66)=3.358, P=0.0408, oEPSCs:HVC_RA_ vs HVC_X_ P=0.0029, HVC_RA_ vs HVC_Av_ P=0.0157); **C**) oPSCs rise latency for cells where both oEPSC and oIPSC were measured (2W ANOVA, oEPSC vs. oIPSC F(1,23)=41.60, P<0.001, HVC_X_ P<0.001, HVC_Av_ P<0.001); **D**) relative delay of oIPSC compared to oEPSC across the cells reported in C) (Kruskal-Wallis test, H=8.809, P=0.0122).

### Novel connection from mMAN to Av and selective monosynaptic connections with HVC

To better define the specificity of regional DVR projections we next mapped synaptic connectivity of two other HVC afferents, mMAN and Av. The DVR is comprised of three large pallial fields separated by distinct lamina, the nidopallium, mesopallium, and the arcopallium. HVC, NIf, and mMAN reside in the nidopallium and Av is in the mesopallium. HVC and NIf are in caudal regions of the DVR’s nidopallium while mMAN is located at anterior portions of the DVR, some 4-5mm from HVC. Av, on the other hand is only ∼1.5 mm from HVC but is in a distinct pallial field.

Cell type specific synaptic connectivity between nidopallial regions and between mesopallial and nidopallial regions are poorly understood; however, the connectivity of mMAN and Av are better described than other portions of these pallial fields. mMAN receives synaptic input from the thalamic nucleus DMP and is part of a vocal motor DVR-thalamo-DVR loop (mMAN > HVC > RA > DMP > mMAN), a circuit architecture similar to premotor and motor control-related cortico-thalamo-cortical loops in mammals. Although the synaptic connectivity between mMAN and HVC has not been well studied, lesions of mMAN in juvenile zebra finches result in song imitation deficits and lesions in adult Bengalese finches increase song syntax variability, suggesting a role in selection and planning of vocal motor sequencing^41,65^. Av is one of the more recently described song related regions and unlike Uva, NIf, and mMAN, it is reciprocally connected to HVC. The role of Av is still poorly explored, but it is understood to be embedded in a higher order auditory portion of the DVR and thought to be an important node in the circuit comparing efferent copies of motor commands from HVC to auditory feedback ^35,39^.

While evaluating connectivity between HVC, mMAN, and Av we discovered that mMAN also provides a strong projection to Av (**Fig. 4**). Approximately half of the identified projection neurons in mMAN project to Av and our tracing experiments reveal that there are only partially overlapping classes of projection neurons in mMAN: mMAN_HVC_ PNs, mMAN_Av_ PNs, and mMAN_HVC/AV_ PNs. This newly identified pathway appears positioned to provide the auditory system with information about song motor commands via a DVR-thalamo-DVR loop (RA > DMP > mMAN > Av). It also opens the potential for synaptic loops relaying through HVC between these regions.

**Fig. 4:**
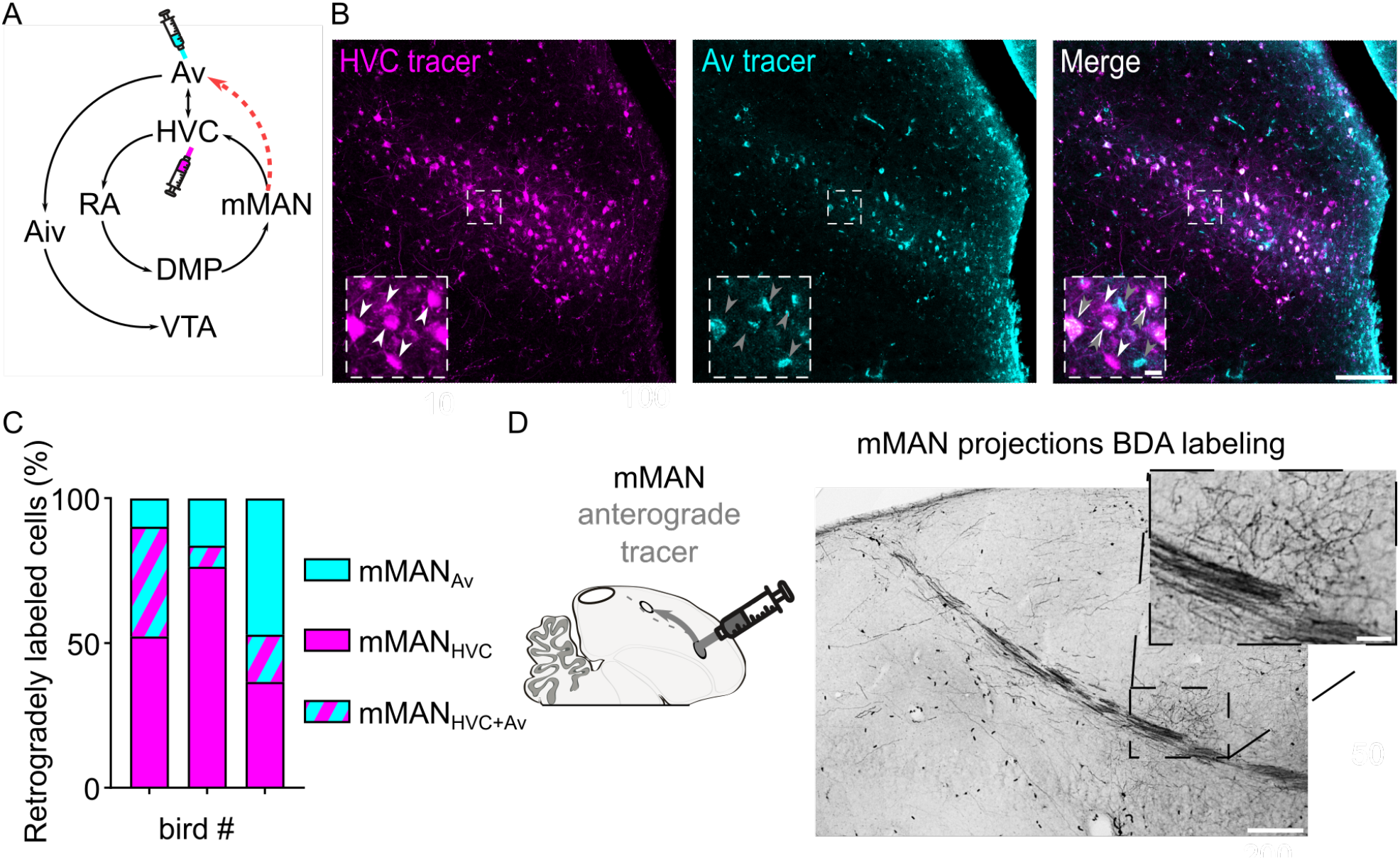
Identification of mMAN neuronal subpopulations projecting to Av and to HVC. **A**) Schematic of the known mMAN afferent and efferent circuitry, together with the proposed new connection towards Av (red dashed arrow), and schematized injection of retrograde tracers in HVC and Av. **B**) sample images of retrogradely labeled mMAN cells projecting to HVC (magenta) or to Av (cyan), and merged image. Insets report magnified selection (dashed square box in the images) illustrating the potential 3 subpopulations identified: mMAN_HVC_ (white arrowheads), mMAN_Av_ (grey arrowheads) and mMAN_HVC+Av_ (overimposed white and grey arrowheads) (scalebars: 100µm, inset: 10µm). **C**) Bar chart displaying the number of retrogradely mMAN_HVC_, mMAN_Av_, and mMAN_HVC+Av_ labeled cells, in 3 birds (averaged across hemispheres, 2-6 slices/bird). **D**) BDA labeling of anterograde projection to Av from mMAN, inset magnifies the terminal field in Av (scalebars: 200µm, inset: 50µm).

Brain regions nearby mMAN do not send projections to HVC and we were able to achieve robust expression of eGtACR1 in mMAN (**Fig. 5A**), allowing us to selectively examine the synaptic connectivity between mMAN and HVC-PNs. Optical stimulation of mMAN axon terminals in brain slices revealed glutamatergic oEPSCs in all retrogradely identified projection neuron classes, albeit with significantly lower success rate compared to what we observed when stimulating NIf terminals (48.15% probability; mMAN vs. NIf oEPSC probability: Fisher’s exact test *P*<0.001). In addition, we found that oEPSCs were more likely evoked in recordings from HVC_X_ and HVC_Av_ than from HVC_RA_ neurons (**Fig. 5B, Suppl. Fig. 3**).

**Fig. 5:**
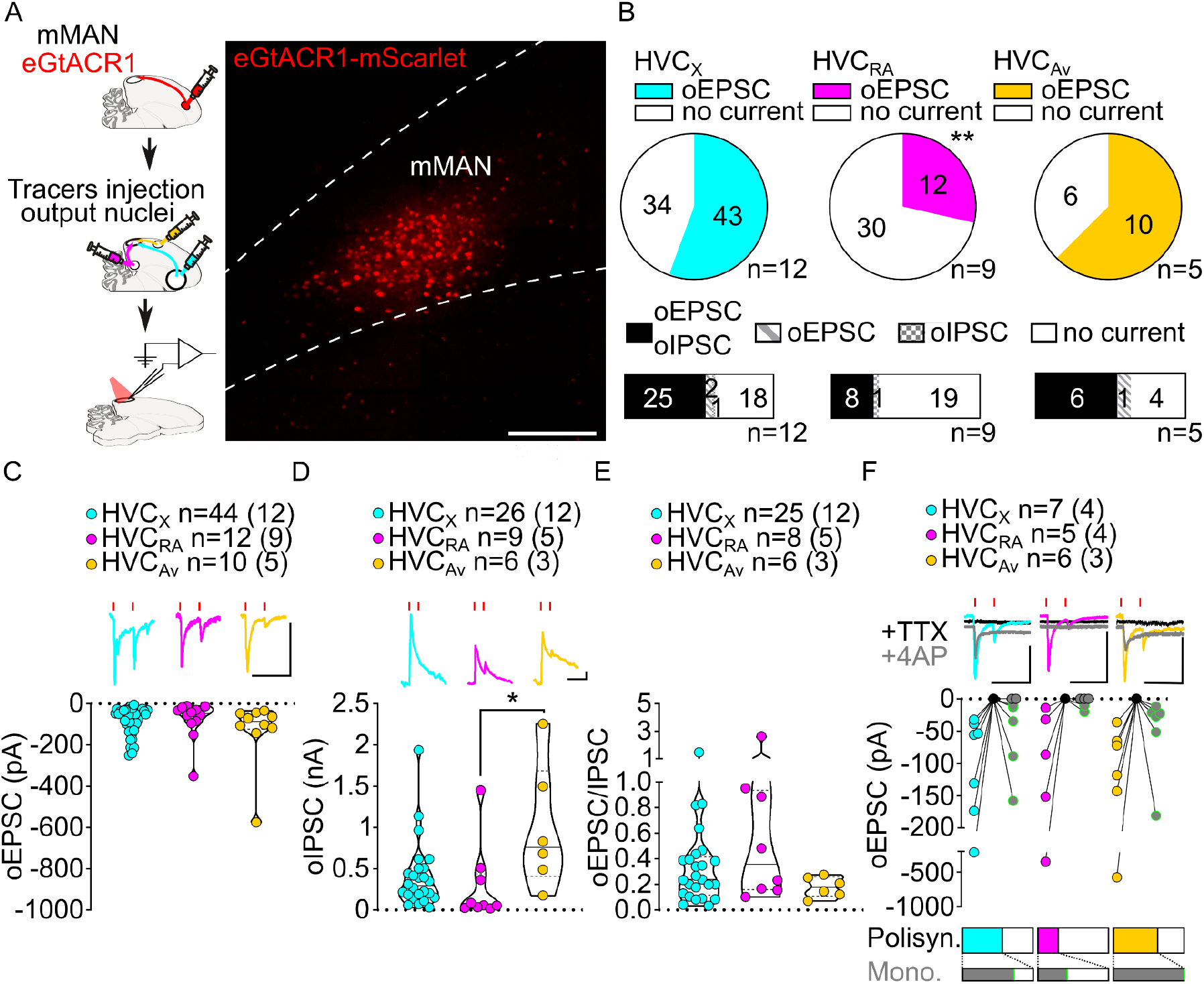
mMAN inputs elicit polysynaptic oIPSCs and oEPSCs in all three HVC-PN classes and monosynaptically excite HVC_X_ and HVC_Av_ neurons. **A**) (left) Schematic of the experimental timeline, illustrating injection of AAV-eGtACR1 in MMAN, followed by retrograde tracer injections in HVC efferent areas and whole-cell patch-clamp recording in acute brain slices; (right) sample image of eGtACR1-mScarlet expression in MMAN (scalebar 200µm). **B**) (top) pie charts representing the likelihood of observing oEPSCs in HVC_X_ (cyan), HVC_RA_ (magenta) or HVC_Av_ (yellow) (Fisher’s exact test, P=0.0088). Numbers in the pie fragments represent the number of cells in which current (colored) or no current (white) was found. Numbers next to the pie charts represent the number of animals from which the data is obtained. (bottom) bar chart representing the number of cells where both oEPSCs and oIPSCs could be elicited (black), only oEPSCs but no oIPSCs (grey lines), only oIPSCs but no oEPSCs (grey checkers), or neither (white), for subsets of cells from the HVC-PN classes’ pie charts aligned above (Fisher’s exact test, P=0.1054). **C**) violin and scatter plot and sample traces reporting average measured oEPSC amplitude of each cell, by cell class (Kruskal-Wallis test H(2)=2.659, P=0.2646; n= cells (animals); red lines represent light stimuli, 1ms; scalebars, 100ms, 100pA). **D**) violin and scatter plot and sample traces of average measured oIPSC amplitude of each cell, by cell class (H(2)=8.598, P=0.0136, HVC_RA_ vs. HVC_Av_ P=0.0102). **E**) violin and scatter plot of the ratio of oEPSC and oIPSC peak amplitude of each cell where both are measured and ≠0, per cell class (H(2)=2.443, P=0.2759). **F**) (top) sample traces and plot representing the amplitude of post-synaptic currents evoked by lightly-driven release of neurotransmitter from MMAN axonal terminals in HVC; oEPSCs amplitudes are reported before (HVC_X_ cyan, HVC_RA_ magenta, HVC_Av_ yellow) and after bath application of TTX (black) and 4AP (grey, green outline indicates polysynaptic oEPSC, see methods), (n= cells (animals); blue lines represent light stimuli, 1ms; scalebars, 100ms, 100pA) (bottom) bar charts representing the likelihood of observing polysynaptic oEPSCs in HVC_X_ (cyan), HVC_RA_ (magenta) or HVC_Av_ (yellow) (data from panel B), and (grey) likelihood of a subset of the corresponding oEPSCs to be monosynaptic.

Optogenetic stimulation of mMAN axon terminals also revealed GABAergic currents in all three HVC-PN classes (48.24% probability). Similar to NIf and Uva inputs, oIPSCs elicited by mMAN axons were most often found in cells where oEPSCs were also observed (oIPSC contingent with oEPSC: HVC_X_: 93.5%, HVC_RA_: 96.4%, HVC_Av_: 90.9%; **Fig. 5B**).

While oEPSCs amplitudes across HVC-PN classes were statistically indistinguishable (**Fig. 5C)**, we found that oIPSCs in HVC_Av_ had significantly higher amplitude than those evoked in HVC_RA_ neurons (**Fig. 5D, Suppl. Fig. 3**). Despite this difference, when computing the E/I ratio we didn’t see any significant difference across HVC-PNs, and like the other inputs examined, GABAergic currents had higher amplitude than glutamatergic currents (**Fig. 5E, Suppl. Fig. 3**).

We next asked whether mMAN monosynaptically contacted any class of HVC-PNs. We found evidence that mMAN monosynaptically contacts all three classes of HVC PNs. The most reliable monosynaptic transmission is between mMAN and HVC_Av_ neurons and HVC_X_ neurons (HVC_RA_: 2/5, HVC_X_: 5/7, HVC_Av_: 6/6, **Fig. 5F**). We did not observe monosynaptic oIPSCs (HVC_X_: 0/5, HVC_RA_, 0/5, HVC_Av_: 0/6, data not shown), and confirmed that mMAN neurons projecting to HVC are glutamatergic using in-situ hybridization combined with retrograde tracing from injection of tracer in HVC (**Suppl. Fig. 3**). Together, this data indicates that mMAN predominantly provides robust monosynaptic excitatory input to HVC_Av_ and HVC_X_ neurons. HVC_Av_ neurons only comprise ∼3% of the neurons in HVC^27^. Therefore, finding that 100% of the HVC_Av_ neurons tested receive monosynaptic input from mMAN suggests that this is likely to be a highly selective synaptic target of mMAN inputs to HVC. Given that we have now also identified a projection from mMAN directly to Av, this supports a model in which mMAN is providing Av song related signals directly and indirectly via projections onto HVC_Av_ neurons.

**Suppl. Fig 3 (related to Fig. 5).**
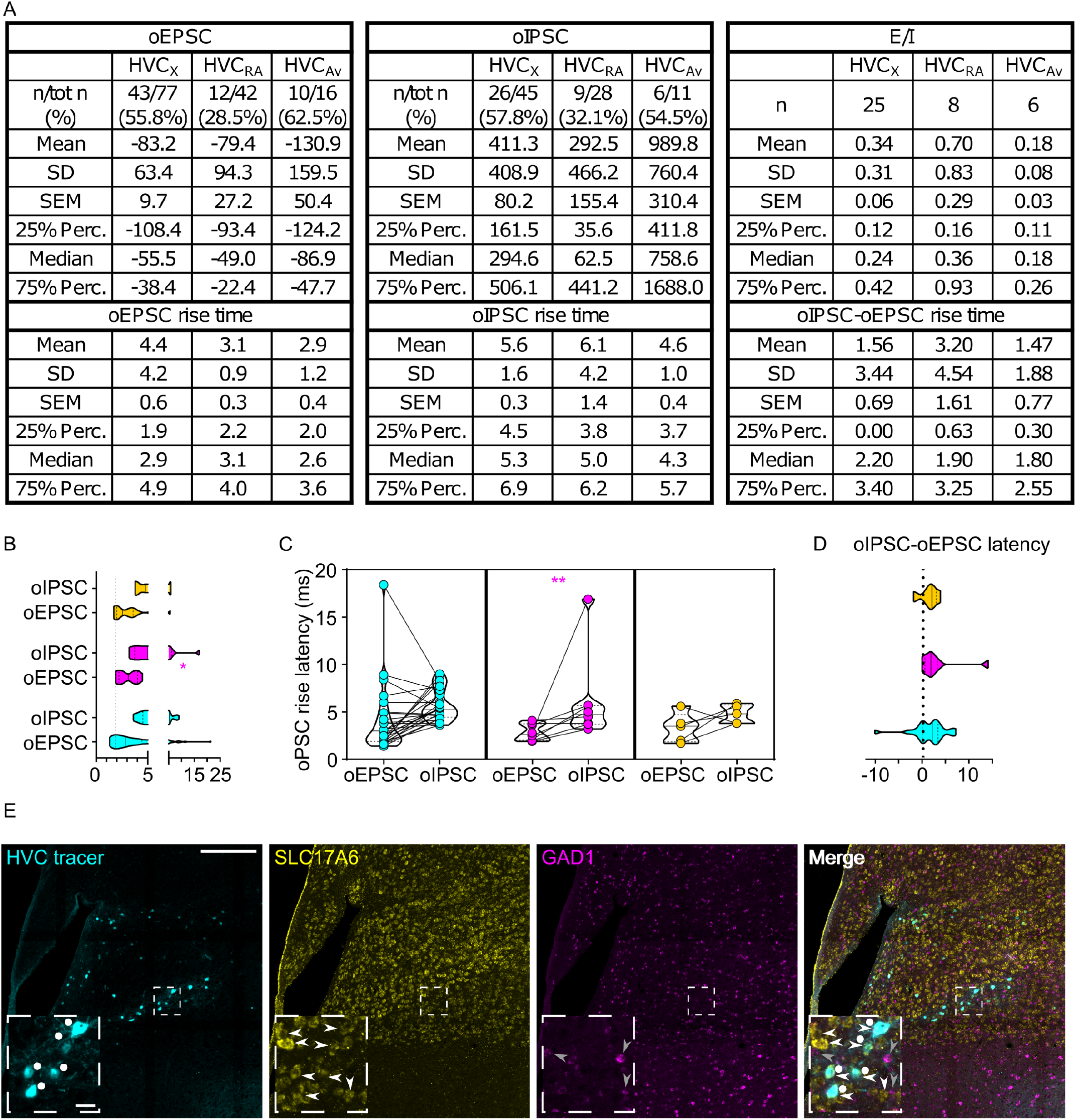
**A**) Table reporting descriptive statistics relative to oEPSCs and oIPSCs as reported in Fig. 5. **B)** violin and scatter plots representing oPSC rise latencies reported in fig. 5 (gray line marks the minimal latency (1_st_ quartile) to oPSC rise across the cell types, for ease of comparison (Mixed-effects analysis, oEPSC vs. oIPSC F(1,35)=8.439, P=0.0063, HVC_RA_ P=0.0198; HVC_PN_ F(2,64)=0.7545, P=0.4744); **C**) oPSCs rise latency for cells where both oEPSC and oIPSC were measured (2W ANOVA, oEPSC vs. oIPSC F(1,35)=8.294, P=0.0067, HVC_RA_ P=0.0470); **D**) relative delay of oIPSC compared to oEPSC across the cells reported in C) (Kruskal-Wallis test, H=0.2028, P=0.9036). **E**) Sample images reporting retrogradely labeled HVC-projecting neurons in mMAN (cyan, white circles) together with in-situ labeling of glutamatergic (SLC17A6, yellow, white arrowheads) and GABAergic (GAD1, magenta, gray arrowheads) markers transcripts (scalebar 200µm, inset 20µm).

Lastly, we mapped the synaptic inputs from Av onto HVC-PNs. **6A**). Avalanche lacks clear anatomical boundaries, and our viral expression did spread into surrounding brain regions, but only brain hemispheres in which we could verify a lack of expression in NIf were included in this study (**Fig. 6A)**. Whole-cell patch-clamp recordings from identified HVC-PN classes in brain slices revealed glutamatergic oEPSCs in all 3 cell classes, yet most of the cells were not excited by Av terminals (39.18% probability; Av vs. NIf oEPSC probability: Fisher’s exact test *P* < 0.001). However, the probability of evoking oEPSCs was similar among HVC-PN classes (**Fig. 6B, Suppl. Fig. 4**). We tested the presence of oIPSCs in a subset of cells and found GABAergic currents in all three projection subtypes (43.24% probability, **Suppl. Fig. 4**). As in other pathways already described, we observed oIPSCs mainly in HVC-PNs also showing oEPSCs (oIPSC contingent with oEPSC: HVC_X_ 100.0% HVC_RA_ 90.0%, HVC_Av_ 84.6%; **Fig. 6B**).

**Fig. 6:**
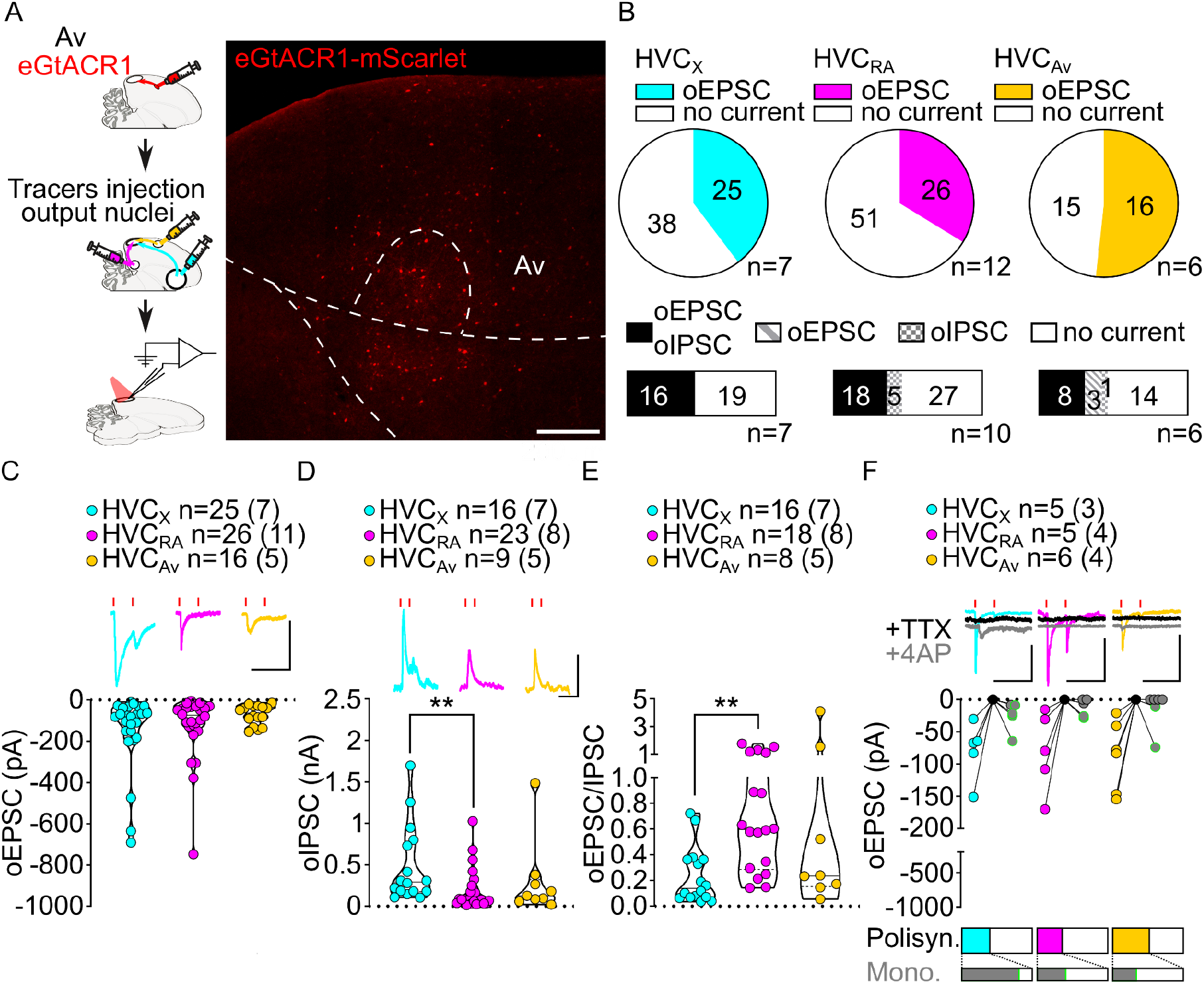
Av inputs elicit polysynaptic oIPSCs and oEPSCs in all three HVC-PN classes and monosynaptically excite mainly HVC_X_ neurons. **A**) (left) Schematic of the experimental timeline, illustrating injection of AAV-eGtACR1 in Av, followed by retrograde tracer injections in HVC efferent areas and whole-cell patch-clamp recording in acute brain slices; (right) sample image of eGtACR1-mScarlet expression in Av (scalebar 200µm). B) (top) pie charts representing the likelihood of observing oEPSCs in HVC_X_ (cyan), HVC_RA_ (magenta) or HVC_Av_ (yellow) (Fisher’s exact test, P=0.2243). Numbers in the pie fragments represent the number of cells in which current (colored) or no current (white) was found. Numbers next to the pie charts represent the number of animals from which the data is obtained. (bottom) bar chart representing the number of cells where both oEPSCs and oIPSCs could be elicited (black), only oEPSCs but no oIPSCs (grey lines), only oIPSCs but no oEPSCs (grey checkers), or neither (white), for subsets of cells from the HVC-PN classes’ pie charts aligned above (Fisher’s exact test, P=0.0550). **C**) violin and scatter plot and sample traces reporting average measured oEPSC amplitude of each cell, by cell class (Kruskal-Wallis test H(2)=1.103, P=0.5760; n= cells (animals); red lines represent light stimuli, 1ms; scalebars, 100ms, 100pA). **D**) violin and scatter plot and sample traces of average measured oIPSC amplitude of each cell, by cell class (H(2)=10.09, P=0.0064, HVC_X_ vs. HVC_RA_ P=0.0047). **E**) violin and scatter plot of the ratio of oEPSC and oIPSC peak amplitude of each cell where both are measured and ≠0, per cell class (H(2)=10.64, P=0.0049, HVC_X_ vs. HVC_RA_ P=0.0033). **F**) (top) sample traces and plot representing the amplitude of post-synaptic currents evoked by lightly-driven release of neurotransmitter from Av axonal terminals in HVC; oEPSCs amplitudes are reported before (HVC_X_ cyan, HVC_RA_ magenta, HVC_Av_ yellow) and after bath application of TTX (black) and 4AP (grey, green outline indicates polysynaptic oEPSC, see methods), (n= cells (animals); blue lines represent light stimuli, 1ms; scalebars, 100ms, 100pA) (bottom) bar charts representing the likelihood of observing polysynaptic oEPSCs in HVC_X_ (cyan), HVC_RA_ (magenta) or HVC_Av_ (yellow) (data from panel B), and (grey) likelihood of a subset of the corresponding oEPSCs to be monosynaptic.

While oEPSCs amplitudes were broadly similar among HVC-PN classes (**Fig. 6C**), oIPSCs had significantly smaller amplitude in HVC_RA_ compared to those evoked in HVC_X_ neurons (**Fig. 6D, Suppl. Fig. 4**). Consistently, this was also reflected in a significantly higher E/I ratio in HVC_RA_ compared to HVC_X_. Similar to the other afferents, Av axon terminal stimulation elicited stronger GABAergic currents than glutamatergic currents across HVC-PN classes (**Fig. 6E, Suppl. Fig. 4**).

When examining which HVC-PN neuron classes received monosynaptic input from Av, we consistently observed monosynaptic transmission onto HVC_X_ neurons (HVC_X_ 4/5 cells monosynaptically contacted). Monosynaptic transmission onto HVC_RA_ and HVC_Av_ was less common (HVC_RA_ 2/5, HVC_Av_ 2/6; **Fig. 6F**). Previous reports indicated that Av neurons projecting to HVC are excitatory^27^. Consistent with this, we didn’t observe any monosynaptic oIPSC across HVC-PNs (monosynaptic oIPSCs: HVC_X_ 0/4, HVC_RA_ 0/3, HVC_Av_ 0/3, data not shown) and further confirmed that Av_HVC_ neurons are glutamatergic by in-situ hybridization labeling of retrogradely identified neurons (**Suppl. Fig. 4**). Our data shows that Av provides monosynaptic excitatory input to HVC_X_ neurons with high frequency and makes monosynaptic connections with HVC_RA_ and HVC_Av_ neurons less frequently.

Taken together, this research provides a rich dataset mapping the polysynaptic and monosynaptic connectivity of the input and output pathways of HVC - the best studied portions of the avian song circuitry – as well as a newly discovered pathway between DVR circuits that are important for vocal learning (**Fig. 7**).

**Suppl. Fig. 4 (related to Fig. 6).**
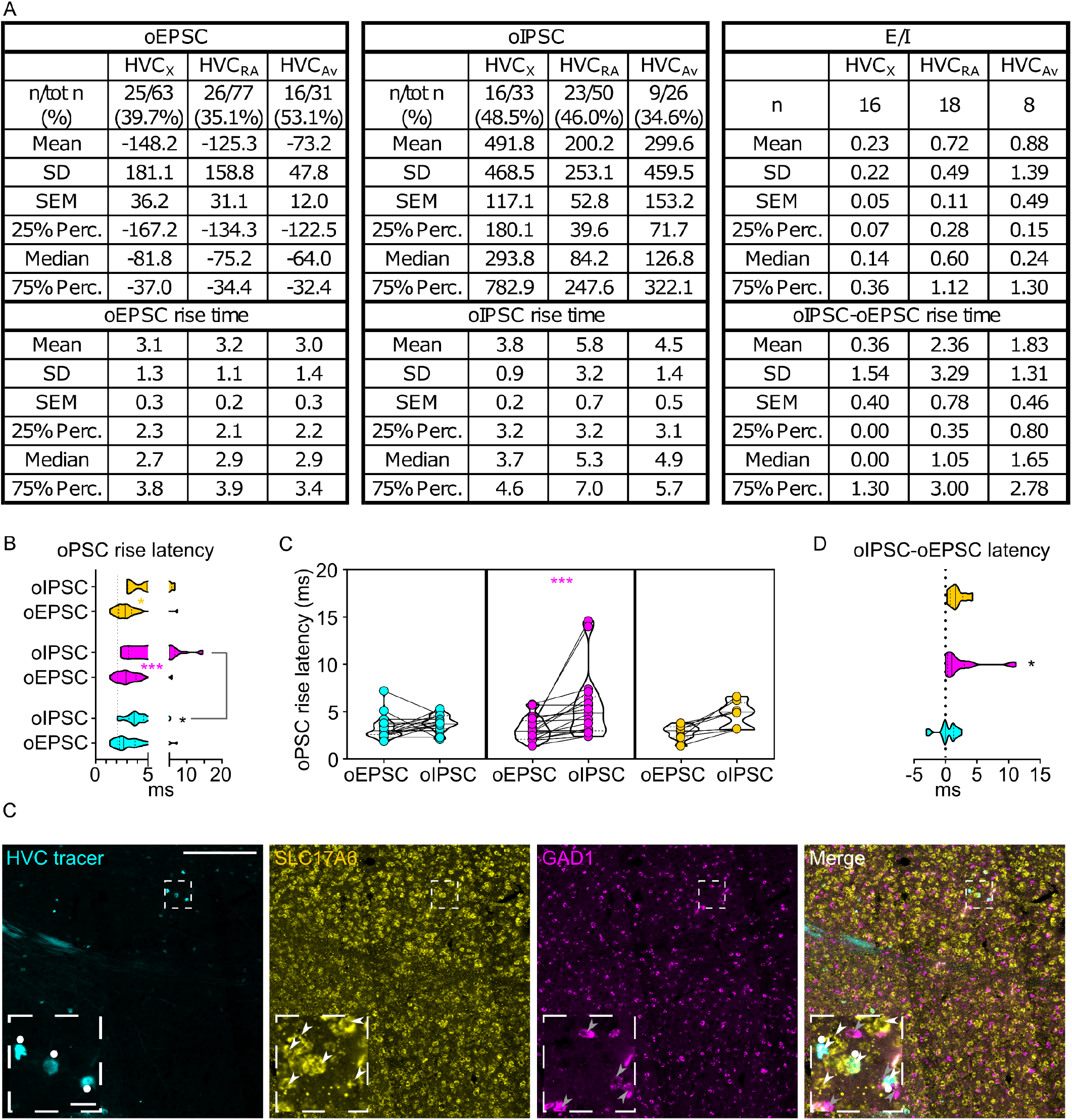
**A**) Table reporting descriptive statistics relative to oEPSCs and oIPSCs as reported in Fig. 6. **B)** violin and scatter plots representing oPSC rise latencies reported in fig. 6 (gray line marks the minimal latency (1_st_ quartile) to oPSC rise across the cell types, for ease of comparison (Mixed-effects analysis, oEPSC vs. oIPSC F(1,39)=21.78, P<0.001, HVC_RA_ P<0.001, HVC_Av_ P=0.0377; HVC_PN_ F(2,70)=2.764, P=0.0699); **C**) oPSCs rise latency for cells where both oEPSC and oIPSC were measured (2W ANOVA, oEPSC vs. oIPSC F(1,39)=14.26, P<0.001, HVC_RA_ P<0.001); **D**) relative delay of oIPSC compared to oEPSC across the cells reported in C) (Kruskal-Wallis test, H=6.403, P=0.0407). **E**) Sample images reporting retrogradely labeled HVC-projecting neurons in Av (cyan, white circles) together with in-situ labeling of glutamatergic (SLC17A6, yellow, white arrowheads) and GABAergic (GAD1, magenta, gray arrowheads) markers transcripts (scalebar 200µm, inset 20µm).

**Fig. 7:**
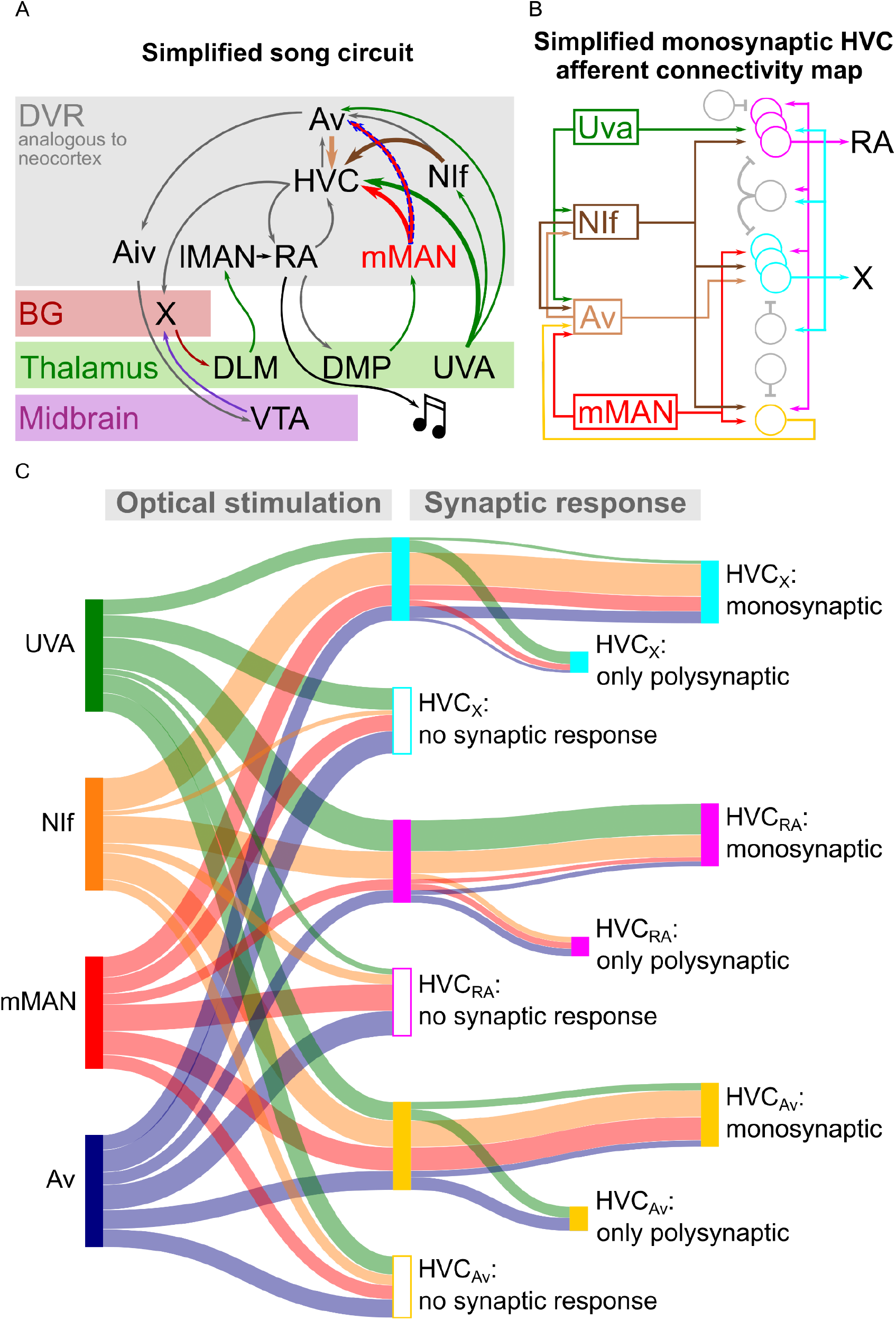
Song circuitry summary and monosynaptic connectivity map of afferents to HVC-PNs. **A**) Schematic representing a simplified version of the long-range connectivity map for song. Colors represent the anatomical location of each projection: gray: DVR, red: basal ganglia, green: thalamus, purple: midbrain. HVC afferents described in this manuscript are highlighted by large arrows. The novel mMAN-Av projection is highlighted by a dashed outline. **B**) Schematic of the HVC afferent connectivity map resulting from the present work, complemented with projections between HVC afferent areas (on the left) and between HVC projection neurons as per previous reports (on the right). For conceptualization purposes, afferent connectivity to HVC-PNs is shown only when the rate of monosynaptic connectivity reaches 50% of neurons examined: NIf monosynaptically contacts all 3 HVC-PNs, while Uva is preferentially monosynaptically connected to HVC_RA_, mMAN to HVC_X_ and HVC_Av_, and Av to HVC_X_. **C**) Sankey diagram displaying the prevalence of connectivity for each input and cell subtype combination, based on polysynaptic and monosynaptic connectivity rates described in figures 2-3 and 5-6.

## Discussion

Speech and language are controlled by interconnected neocortical and subcortical networks, including dorsal and ventral cortical pathways for articulation and lexical processing, and pathways looping through the thalamus, basal ganglia, and cerebellum^6,66-68^. The synaptic connectivity between specific populations of neurons among these circuits is going to be difficult to map. Yet, this will ultimately be necessary for understanding the neuroanatomical basis and circuit computations associated with speech and language. Although the avian DVR and mammalian neocortex have distinct developmental origins (ventral and dorsal pallium, respectively), HVC PNs neurons have similar gene expression and connectivity patterns to the intratelencephalic neurons described in layers 2-6 of the mammalian neocortex^32^, suggesting evolutionary convergence in circuit computations needed for complex behaviors like vocal imitation. Thus, there is much to be gained by mapping the connectivity between the specific populations of neurons known to be essential in the sensory and sensorimotor process of song imitation.

Using large-scale, multi-pathway, and long-range functional synaptic circuit mapping, we report cell type specific synaptic connectivity across forebrain sensorimotor networks necessary for learning and producing learned vocalizations. The songbird motor nucleus HVC is crucial for song production. Its partial or complete lesion results in song disruption and permanent loss, respectively^28,69,70^. Electrical stimulation of HVC can halt song^71-73^. Further, HVC is essential in juvenile song learning ^27,36,46,74,75^. We show that long-range sensory pathways to HVC are excitatory and contact HVC intratelencephalic PNs. We find that these connections are not stochastic but are instead strongly biased to make synapses with premotor (HVC_RA_), auditory (HVC_AV_), or basal ganglia (HVC_X_) projection pathways. Lastly, we find a remarkable correspondence in the probability of finding oEPSCs and oIPSCs in the same postsynaptic neurons, indicative of highly interconnected cell assemblies at postsynaptic targets of long-range connections within the DVR.

In addition to finding widespread polysynaptic transmission upon stimulation of any of the afferents, we endeavored to reveal the wiring diagram of HVC inputs by isolating the monosynaptic components of these currents (**Fig. 7**). We found surprisingly specific and compartmentalized monosynaptic neurotransmission between each input and HVC-PN classes. NIf terminals monosynaptically contacted all three HVC-PN classes. Uva predominantly makes monosynaptic connections with HVC_RA_ neurons, while mMAN most frequently makes monosynaptic connections with HVC_Av_ and HVC_X_ PNs but appears to only sparsely project to HVC_RA_ neurons. Lastly, Av predominantly makes monosynaptic connections with HVC_X_ neurons and connects less frequently with HVC_RA_ and HVC_Av_ neurons. When combined with our discovery of a new pathway from mMAN to Av, our data draws a scenario of multiple intermingled loops coexisting across HVC input and output circuits.

Thalamic inputs from Uva may directly affect the song motor pathway through monosynaptic projections onto HVC_RA_ neurons. The potential relevance of this specific connection has been recently studied^29,45^. Additionally, the seldom yet significant direct inputs to HVC_Av_ neurons we identify are interesting considering the proposed role of HVC_Av_ neurons in providing a forward model of song to the auditory system^27^. Uva also is known to project to both NIf and Av. Uva’s monosynaptic connections to classes of HVC PNs may therefore directly integrate with propagation of forward models of song timing to the auditory system, particularly during learning of syllable transitions^27^.

mMAN has recently been found to contribute to the ordering of song syllables^76^. This could be attributable to the strong connections identified here between mMAN and HVC_Av_ PNs and from mMAN directly to Av. HVC_Av_ neurons are important for learning the timing of elements within song. Lesions of HVC_Av_ neurons in juvenile birds disrupts learning of syllable syntax yet does not have a strong effect on learning the spectral features of syllables^27^. Further, lesions block deafening-induced degradation of songs’ temporal features and recovery of song timing following song-contingent feedback perturbations^27^. Thus, our analysis of synaptic connectivity provides important anatomical connections for a circuit critical for temporal ordering of vocalizations during learning and in adulthood.

mMAN and Av’s preferential connection to HVC_X_ PNs may create a bridge linking the thalamus, the auditory system, and the basal ganglia circuits important for song plasticity. mMAN receives projections from the nucleus DMP of the thalamus, which itself receives projections from RA. mMAN is therefore positioned to receive a copy of descending vocal motor commands transmitted through DMP while Av may relay integrated information about auditory feedback and their correspondence with motor commands for song that it receives from mMAN and from HVC_Av_ neurons. Our findings place mMAN and Av in an ideal position to relay those signals to Area X, potentially contributing to song plasticity.

In the data shown in Figs. **2**, **3, 5-6** we examined the probability and strength of oPSCs within each HVC-PNs class. We found that projection neurons in NIf, Uva, mMAN and Av consistently establish polysynaptic contact with all three HVC-PN classes. We reliably observed both glutamatergic and GABAergic polysynaptic neurotransmission upon afferent axonal terminal stimulation. Organizing our data by HVC PN types (**Suppl. Fig. 5**) reveals a bias in afferent inputs onto these different neuronal pathways. We find that HVC_X_ and HVC_AV_ neurons are most frequently excited by NIf afferents, while HVC_RA_ are most frequently excited by Uva, and unlikely to be excited by mMAN terminal stimulation. HVC_X_ and HVC_Av_ neurons are most strongly driven by NIf, while HVC_RA_ neurons receive relatively uniform amplitude oEPSCs from all 4 input pathways examined. Lastly, stimulation of NIf and mMAN terminals resulted in higher amplitude inhibition onto HVC_Av_ neurons than upon Uva or Av terminals stimulation (**Suppl. Fig. 5**).

This data points towards the existence of a rich interneuronal inhibitory network that is recruited by stimulating HVC’s afferents, evoking GABAergic transmission onto all HVC-PN classes. This is in line with previous reports that found HVC-PNs activity to be tightly regulated by interneurons^77-79^. Interestingly, across all inputs studied, the likelihood of finding oIPSCs was tightly dependent on finding oEPSCs in the same cell. Previous studies have indicated that HVC PNs are disynaptically inhibited by other HVC-PNs, a feature of synaptic connectivity that is likely fundamental to the propagation of excitation in the network during song production and in the song learning process ^77,79^. When afferents cause excitation of HVC PNs this would therefore cause an immediate rise in the levels of disynaptic inhibition in the surrounding HVC PNs. Similar feedback inhibition dynamics have been extensively reported in cortical circuits, where this mechanism serves a crucial role of controlling the excitatory-inhibitory balance of local principal neurons^80-83^. Afferents that excite HVC PNs may also contact neighboring interneurons. A similar feedforward inhibition has also been reported in the cortex, where both pyramidal neurons and interneurons projecting to them are recruited by long-range pathways ^84,85^. While feedback inhibition is poised to monitor and modulate the output of local principal neurons, feedforward inhibition is proposed to work as a coincidence detector improving the sensitivity of the network by filtering asynchronous inputs^80,86-90^

**Suppl. Fig. 5:**
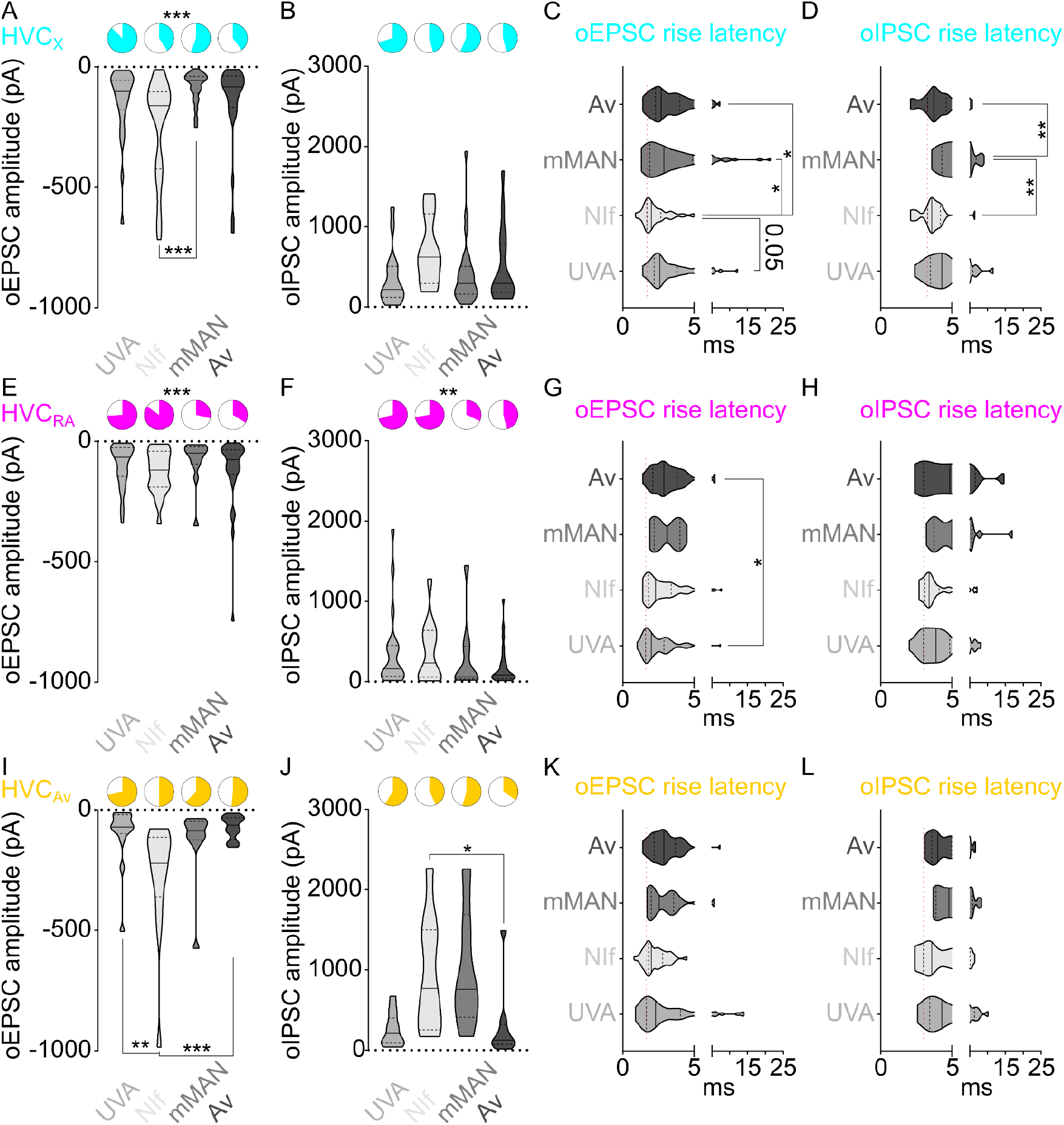
Cross-comparison of afferent polysynaptic connectivity within cell types (data from Fig. 2,3,5,6). **A**) Pie charts representing the likelihood of evoking oEPSCs in HVC_X_, and violin plots representing the oEPSCs amplitude, when stimulating afferents from NIf, UVA, mMAN or Av (probability: Fisher’s exact test, P<0.001; amplitude: Kruskal-Wallis test, H(3)=17.18, P<0.001, NIf vs. UVA P=0.1695, NIf vs. mMAN, P<0.001, NIf vs. Av P=0.0993). **B**) Same as A) for oIPSCs (probability: Fisher’s exact test, P=0.2828; amplitude: Kruskal-Wallis test, H(3)=5.008, P=0.1712). **C**) Violin plots reporting the oEPSC rise latency in HVCX upon stimulation of the 4 afferents (Kruskal-Wallis test, H(3)=11.91, P=0.0077, NIf vs. UVA P=0.0534, NIf vs. mMAN, P=0.0119, NIf vs. Av P=0.0345). **D**) same as C) for oIPSCs (Kruskal-Wallis test, H(3)=19.23, P<0.001, NIf vs. UVA P=0.5428, NIf vs. mMAN, P=0.0016, NIf vs. Av P>0.9999, mMAN vs. Av P=0.0028). **E-H**) Same as A-D) but for HVC_RA_: E) (probability: Fisher’s exact test, P<0.001; amplitude: Kruskal-Wallis test, H(3)=3.594, P=0.3088). F) (probability: Fisher’s exact test, P=0.0075; amplitude: Kruskal-Wallis test, H(3)=2.132, P=0.5454). G) (Kruskal-Wallis test, H(3)=12.44, P=0.0060, NIf vs. UVA P>0.999, NIf vs. mMAN, P=0.6258, NIf vs. Av P=0.3175, UVA vs. Av P=0.0128). H) (Kruskal-Wallis test, H(3)=6.622, P=0.0850). **I-L**) Same as A-D) but for HVC_Av_: I) (probability: Fisher’s exact test, P=0.3150; amplitude: Kruskal-Wallis test, H(3)=18.46, P<0.001, NIf vs. UVA P=0.0016, NIf vs. mMAN, P=0.0929, NIf vs. Av P=0.0010). J) (probability: Fisher’s exact test, P=0.1869; amplitude: Kruskal-Wallis test, H(3)=12.52, P=0.0058, NIf vs. UVA P=0.1043, NIf vs. mMAN, P>0.999, NIf vs. Av P=0.0405). K) (Kruskal-Wallis test, H(3)=6.441, P=0.0920). L) (Kruskal-Wallis test, H(3)=7.653, P=0.0538).

Distinguishing among these potential feedback and feedforward circuit motifs, as well as fully characterizing the role of afferents for HVC local circuit dynamics, will require monosynaptic connectivity mapping of afferents onto HVC interneurons. The ability to do this systematically in HVC is limited by the lack of genetic methods for labeling interneurons^91^. Colquitt et al.^32^ have recently identified multiple GABAergic cell subtypes in HVC, and a molecular handle is needed to experimentally approach the study of their synaptic connectivity. Nonetheless, our data suggests the existence of compartmentalized neuronal ensembles including both PNs and interneurons projecting to those PNs.

The synaptic organization of excitatory-inhibitory connections in HVC seem to allow the integration of afferent inputs within hyper-local networks or modules. This type of connectivity may make the circuit more robust to perturbations^92^. A potential caveat is that we performed all our recordings in 230µm thick sagittal brain slices and it is thought that HVC-PNs exhibit sequential activity that is organized in the rostrocaudal axis, while interneurons are speculated to be coordinated in the medio-lateral axis^93^. Our resection of this mediolateral organization may account for the lack of lateral inhibition found in our data. For example, our optogenetic stimulation elicits oIPSCs mostly in the same neurons where oEPSCs are observed, but not in the others, which may have received inhibition from interneurons sitting in adjacent brain slices. Lateral inhibition is normally observed in circuits displaying feedback inhibition, and previous reports indicate that HVC-PNs disynaptically inhibit other HVC-PNs ^77,79^. It is however possible that differences between neocortical and HVC microcircuitry may reduce the significance of lateral inhibition, explaining why we only seldomly observed it.

Our measurements of oPSCs onset delays from the light stimulation offer some insight into the possible wiring of afferents and intra-HVC connectivity. HVC_RA_ neurons displayed the fastest oEPSCs latency upon Uva axon terminal stimulation, consistent with their preferential monosynaptic connectivity. Likewise, HVC_X_ neurons display the fastest latency upon NIf axon terminal stimulation. However, cytoarchitecture, axonal transmission latency and synaptic delays may all play roles in determining this timing and other monosynaptic connections are not readily reflected through polysynaptic oPSCs latency measurements. This highlights the importance of using opsin-assisted monosynaptic mapping rather than timing to identify the details of synaptic connectivity.

While our opsin-assisted circuit mapping provides us with a new level of insight into HVC synaptic circuitry, there are limitations to this research that should be considered. All circuit mapping in this study was carried out in brain slices from adult male zebra finches. Future studies will be needed to examine how this adult connectivity pattern relates to patterns of connectivity in juveniles during sensory or sensorimotor phases of vocal learning and connectivity patterns in female birds. Ex-vivo brain slices have historically been demonstrated to be a reliable window into neuronal and circuit function. However intrinsic activity patterns of some neuronal subtypes are known to change in acute slices^94^. Moreover, the slicing procedure severs both neuromodulatory and neurotransmitter afferents, potentially attenuating or removing synaptic inputs to some circuits^95^. Our axonal terminal optical manipulation recruits all the fibers expressing eGtACR1 under the cone of light of the microscope objective. Therefore, our estimates of postsynaptic responses and likelihood of finding responses are based on the simultaneous release of neurotransmitter from all the afferent terminals expressing eGtACR1, and they should be understood in this context. Future studies will need to employ conditions of minimal stimulation that may further characterize the specific properties of monosynaptic and polysynaptic neurotransmission in the examined inputs. Lastly, the amplitude and likelihood of recording oPSCs directly depends on the expression rate and tropism of our viruses, and this may differ across the 4 afferent pathways studied here. We report low levels of viral expression in Av, which may skew our results. However, by testing oPSCs in all three HVC_PN_ classes in each bird helps ensure that the relative contribution Av afferent inputs onto different classes of HVC neurons is not simply an artifact associated with efficiency of viral transduction.

Although a complete description of HVC circuitry will require the examination of other potential inputs (i.e. RA_HVC_ PNs, NCM, A11 glutamatergic neurons^40,96^) and a characterization of interneuron synaptic connectivity, here we provide a map of the synaptic connections between the 4 best described afferents to HVC and its 3 populations of projection neurons in adult birds. These results provide a view into the synaptic underpinnings of circuits fundamental for the learning and production of vocalizations. Moreover, they reveal essential connectivity patterns that can underlie song motor control and learning of syllable order. This picture of input-output organization spanning sensory, thalamic, and premotor areas may thus offer insights into the circuits for human language learning and production. Lastly, the opsin-assisted circuit mapping strategy we used provides an approach to fully characterize other long range and local circuitry in the songbird brain, which will be critical for understanding and modelling the remarkable vocal imitation abilities of songbirds.

## Materials and Methods

### Animals

Experiments described in this study were conducted using adult male zebra finches (130-500 days post hatch). All procedures were performed in accordance with protocols approved by Animal Care and Use Committee at UT Southwestern Medical Center.

### Viral Vectors

The following adeno-associated viral vectors were used in these experiments: rAAV2/9/CBh-FLP and rAAV-CBh-eGtACR1-mScarlet (IDDRC Neuroconnectivity Core at Baylor College of Medicine or UT Southwestern AAV Viral Vector Core). All viral vectors were aliquoted and stored at -80°C until use.

### Constructs

CBh-eGtACR1-mScarlet was generated by inserting KpnI and AgeI restriction sites flanking EF1a in the pAAV-EF1a-F-FLEX-mNaChBac-T2A-tdTomato (a gift from Dr. Massimo Scanziani; Addgene plasmid # 60658) through pcr cloning. The CBh promoter was isolated from pX330-U6-Chimeric_BB-CBh-hSpCas9 (a gift from Dr. Feng Zhang; Addgene plasmid # 42230) by digestion of the KpnI and AgeI restriction sites and inserted into the KpnI/AgeI restriction sites of the pAAV-EF1a-F-FLEX-mNaChBac-T2A-tdTomato backbone. Next, we isolated mScarlet from pmScarlet_C1 (a gift from Dr. Dorus Gadella; Addgene plasmid # 85042) through pcr amplification, inserting a BamHI restriction site into the 5’ end of the ORF. We digested the amplified fragments with BamHI and BsrGI, inserting the digested fragments into BamHI/BsrGI restriction sites of the pAAV-CBh-F-FLEX-mNaChBac-T2A-tdTomato backbone. eGtACR1 was then amplified from pAAV-CAG-DIO-NLS-mRuby3-IRES-eGtACR1-ST (a gift from Dr. Hillel Adesnik; Addgene plasmid # 109048) with a BmtI restriction site on the 5’ end and a AflII restriction site on the 3’end, avoiding amplification of the soma-targeting sequence in the original plasmid. We then digested the amplified fragment with BmtI and AflII, inserting the fragment into the BmtI/AflII sites of the pAAV-CBh-F-FLEX-mNaChBac-T2A-mScarlet.

For CBh-Flpo, we amplified the FLP ORF from pCAG-Flpo (a gift from Dr. Massimo Scanziani; Addgene plasmid # 60662) with a AgeI restriction site on the 5’ end. We then digested the amplified fragment with AgeI and EcoRI, inserting the fragment into the AgeI/EcoRI sites of the CBh-eGtACR1-mScarlet.

Constructs were verified with Sanger sequencing.

### Stereotaxic Surgery

All surgical procedures were performed under aseptic conditions. Birds were anaesthetized using isoflurane inhalation (0.8-1.5%) and placed in a stereotaxic surgical apparatus. The centers of HVC, NIf, mMAN and RA were identified with electrophysiological recordings, and Area X, Av and Uva were identified using stereotaxic coordinates (approximate stereotaxic coordinates relative to interaural zero and the brain surface: head angle, rostral-caudal, medial-lateral, dorsal-ventral (in mm). HVC: 45°, AP 0, ML ±2.4, DV -0.2-0.6; NIf: 45°, AP 1.75, ML ± 1.75, DV -2.4 -1.8; mMAN: 20°, AP 5.1, ML ±0.6, DV -2.1 -1.6; RA: 70°, AP -1.5, ML ± 2.5, DV -2.4 -1.8; X: 45°, AP 4.6, ML ±1.6, DV -3.3 -2.7; Av: 45°, AP 1.65, ML ± 2.0, DV -0.9; UVA: 20°, AP 2.8, ML ±1.6, DV -4.8 -4.2).

Viral injections were performed using previously described procedures^27,46^. Briefly, a cocktail of adeno-associated viral vectors (1:2 of rAAV-CBh-FLP and rAAV-DIO-CBh-eGtACR1, respectively) were injected into NIf, Uva, mMAN or Av (1nl/s, for a total of 1.5 µl/hemisphere). A minimum of 4 weeks after viral injections fluorophore-conjugated retrograde tracers (Dextran 10,000MW, AlexaFluor 488 and 568, Invitrogen; FastBlue, Polysciences) were injected bilaterally into Area X, Av, and RA. Tracer injections (160nl, 5×32nl, 32nl/s every 30 s) were performed using previously described procedures^27,46,97^.

### *In-vivo* extracellular recordings

To test the functional expression of eGtACR1 in HVC afferent pathways, we performed extracellular recording of HVC activity in anesthetized birds previously injected with rAAV-DIO-CBh-eGtACR1 and rAAV-CBh-FLP (2:1). We performed the recordings under light isoflurane anesthesia (0.8%) with Carbostar carbon electrodes (impedance: 1670 microhms/cm; Kation Scientific). During the recordings, we lowered a 400µm multimodal optical fiber to the brain surface overlaying HVC and delivered light stimulation (470nm, ≈20mW, 1s). Signals were acquired at 10KHz, and filtered (high-pass 300Hz, low-pass 20KHz). We used Spike2 to analyze the spike rate (binned every 10ms) and build a peri-stimulus time histogram to evaluate the effect of light stimulation across trials (5-15 trials/hemisphere). We sampled a minimum of 1 site and a maximum of 3 HVC sites/hemisphere. Birds with weak or no optically evoked responses were excluded from further experiments.

### *Ex vivo* physiology

#### Slice preparation

Zebra finches were deeply anesthetized with isoflurane and decapitated. The brain was removed from the skull and submerged in cold (1– 4°C) oxygenated dissection buffer. Acute sagittal 230 μm brain slices were cut in ice-cold carbogenated (95% O2/5% CO2) solution, containing (in mM) 110 choline chloride, 25 glucose, 25 NaHCO3, 7 MgCl2, 11.6 ascorbic acid, 3.1 sodium pyruvate, 2.5 KCl, 1.25 NaH2PO4, 0.5 CaCl2; 320-330 mOsm. Individual slices were incubated in a custom-made holding chamber filled with artificial cerebrospinal fluid (aCSF), containing (in mM): 126 NaCl, 3 KCl, 1.25 NaH_2_PO_4_, 26 NaHCO_3_, 10 D-(+)-glucose, 2 MgSO_4_, 2 CaCl_2_, 310mOsm, pH 7.3–7.4, aerated with a 95% O_2_/5% CO_2_ mix. Slices were incubated at 36 °C for 20 minutes, and then kept at RT for a minimum of 45 minutes before recordings.

#### Slice electrophysiological recording

Slices were constantly perfused in a submersion chamber with 32°C oxygenated normal aCSF. Patch pipettes were pulled to a final resistance of 3-5 MΩ from filamented borosilicate glass on a Sutter P-1000 horizontal puller. HVC-PN classes, as identified by retrograde tracers, were visualized by epifluorescence imaging using a water immersion objective (×40, 0.8 numerical aperture) on an upright Olympus BX51 WI microscope, with video-assisted infrared CCD camera (Q-Imaging Rolera). Data were low-pass filtered at 10 kHz and acquired at 10 kHz with an Axon MultiClamp 700B amplifier and an Axon Digidata 1550B Data Acquisition system under the control of Clampex 10.6 (Molecular Devices).

For voltage clamp whole-cell recordings, the internal solution contained (in mM): 120 cesium methanesulfonate, 10 CsCl, 10 HEPES, 10 EGTA, 5 Creatine Phosphate, 4 ATP-Mg, 0.4 GTP-Na (adjusted to pH 7.3-7.4 with CsOH).

Optically evoked post-synaptic currents (oPSCs)were measured by delivering 2 light pulses (1 ms, spaced 50ms, generated by a CoolLED *p*E300) focused on the sample through the 40X immersion objective. Sweeps were delivered every 10 s. Synaptic responses were monitored while holding the membrane voltage at -70mV (for excitatory oPSCs (oEPSCs)) and +10mV (for inhibitory oPSCs (oIPSCs)). The light stimulation mostly elicited excitatory currents with amplitude in the range of ∼50-600pA. We calibrated the light stimulation intensity to ∼50% of the maximum amplitude. If the amplitude exceeded 600pA, we further reduced the light intensity. In cases with currents below 50pA, we set 20pA as the minimum cut-off current amplitude in order to proceed with experiments. When both oIPSC and oEPSC were measured, the measurement was conducted at the same light stimulation intensity to allow direct comparison. . Access resistance (10–30 MΩ) was monitored throughout the experiment, and for reliability of amplitude measurement recordings where it changed more than 20% were discarded from further analysis. We calculated the paired-pulse ratio (PPR) as the amplitude of the second peak divided by the amplitude of the first peak elicited by the twin stimuli, however due to slow kinetics of eGtACR1 the results would be difficult to interpret, and therefore we are not currently reporting them. The excitatory-inhibitory (E/I) ratio was calculated by dividing the amplitude of the oEPSC at -70mV by the amplitude of the oIPSC at +10mV, while stimulating at identical light intensities. To validate inhibitory and excitatory oPSCs as GABAergic and glutamatergic, respectively, we bath applied the GABAa receptor antagonist SR 95531 hydrobromide (gabazine, µM) while holding the cell at +10mV, or the AMPA receptor antagonist 6,7-Dinitroquinoxaline-2,3-dione (DNQX, µM) while holding the cell at - 70mV. In another subset of cells, once the baseline measures were established, we tested for monosynaptic connectivity. To isolate monosynaptic currents driven by optogenetic stimulation we bath applied 1µM Tetrodotoxin (TTX), followed by 100µM 4-Aminopyridine (4AP) and measured the amplitude of oPSCs returning following 4-AP application. Currents under 5pA were considered not reliable, based on the signal to noise of the recordings, and were assigned value 0 for further analysis. Therefore, average amplitudes and plots relative to oPSCs amplitudes (panels C, D of figures 2,3,5,6) report only cells whose values surpass this 5pA threshold. oPSCs = 0pA were considered to calculate the rate of success in evoking oPSCs and to evaluate whether currents are monosynaptic following TTX and 4AP application. For sake of classification, any current rescued by 4AP application with an amplitude lower than 10pA was considered non-monosynaptic. Birds where no oPSC was recorded in any HVC_PN_ throughout the experimental day were excluded from further analysis.

### Histology

After electrophysiological recordings, the slices were incubated in 4% PFA in PBS. Sections were then washed in PBS, mounted on glass slides with Fluoromount-G (eBioscience, CA, USA), and visualized under an LSM 880 laser-scanning confocal microscope (Carl Zeiss, Germany).

### In-situ hybridization

We used the hairpin chain reaction system from Molecular Instruments as previously described (Ben Tov et al., 2023). Briefly, 5 days after having injected HVC with the retrograde tracer Fast Blue (150nl/HVC, Polysciences) the brain was perfused with 4% paraformaldehyde (PFA), post-fixed at 4 °C for 12 hours and dehydrated in 30% sucrose/4% PFA at 4 °C for 24 hours. The dehydrated brains were sagittally cyrosectioned at 40 μm and collected into ice-cold 4% PFA. Sections were then incubated twice in PBS for 3 minutes, in 5%SDS/PBS for 45 minutes, three times in 2X SSCT for 15 minutes, in Hybridization Buffer for 5 minutes, and finally in 10nM each probe/Hybridization Buffer at 37 °C for 24 hours. The sections were then washed four times in Probe Wash Buffer for 15 minutes at 37 °C, three times in 2X SSCT for 15 minutes at room temperature, then incubated in Amplification Buffer for 30 minutes at room temperature. Each Alexa fluor-conjugated hairpin was denatured in separated PCR tubes at 95 °C for 90 seconds and cooled down to room temperature at 2 °C/s. Sections were incubated in solutions containing 36nM of each hairpin in Amplification Buffer at room temperature for 24 hours. The sections were then washed four times with 2X SSCT for 15 minutes and mounted with mounting medium.

### Experimental Design and Analysis

Electrophysiological data were analyzed with Clampfit (Molecular Devices). All data were tested for normality using the Shapiro-Wilk Test. Parametric and non-parametric statistical tests were used as appropriate. One-way Anova or Kruskal-Wallis test was performed when comparisons were made across more than two conditions. Two-Way Anova or Mixed-effect analysis were used to compare multiple conditions across different groups. Fisher’s exact tests were used to compare the probability of finding optically evoked responses. Statistical significance refers to *P < 0.05, ** P < 0.01, *** P < 0.001.

The Sankey diagram in Fig. 7C was drawn with SankeyMATIC (www.sankeymatic.com) setting ribbons proportional to the percentage of interrogated synaptic connections between each afferent area and HVCPN class displaying excitatory currents or lack thereof. From the hubs reporting existing synaptic connection (center), ribbons are scaled based on the percentage of monosynaptic and polysynaptic connectivity for each combination.

## Author contribution

MT and TFR conceived the project. TFR supervised the research. MT and TFR designed the experiments and wrote the manuscript. MT collected and analyzed the data. ZZ conducted the in-situ hybridization staining. DHA designed and optimized viral tools used in this research. ESM and MZI performed the tracer injections and quantification of mMAN cells. All authors read and commented on the manuscript. Authors declare no competing interests. All data is available in the main text or the supplementary materials.

## Acknowledgments

We thank Drs. David Perkel, Frank Meye, Salvatore Lecca and Manuel Mameli for discussion and comments on an initial version of the manuscript. We thank Andrea Guerrero, Luis Garcia, and Jennifer Holdway for laboratory support. This research was supported by grants from the US National Institutes of Health UF1NS115821 and R01NS108424 to TFR. DA was supported by F99NS124172.

## References

1 Steinmetz, N. A. et al. Neuropixels 2.0: A miniaturized high-density probe for stable, long-term brain recordings. Science 372 (2021). 10.1126/science.abf4588

2 Demas, J. et al. High-speed, cortex-wide volumetric recording of neuroactivity at cellular resolution using light beads microscopy. Nat Methods 18, 1103–1111 (2021). 10.1038/s41592-021-01239-8

3 Manley, J. et al. Simultaneous, cortex-wide dynamics of up to 1 million neurons reveal unbounded scaling of dimensionality with neuron number. Neuron (2024). 10.1016/j.neuron.2024.02.011

4 Consortium, T. M. et al. Functional connectomics spanning multiple areas of mouse visual cortex. bioRxiv, 2021.2007.2028.454025 (2023). 10.1101/2021.07.28.454025

5 Petreanu, L., Mao, T., Sternson, S. M. & Svoboda, K. The subcellular organization of neocortical excitatory connections. Nature 457, 1142–1145 (2009). 10.1038/nature07709

6 Konopka, G. & Roberts, T. F. Insights into the Neural and Genetic Basis of Vocal Communication. Cell 164, 1269–1276 (2016). 10.1016/j.cell.2016.02.039

7 Doupe, A. J. & Kuhl, P. K. Birdsong and human speech: common themes and mechanisms. Annu Rev Neurosci 22, 567–631 (1999).

8 Immelmann, K. in Bird Vocalisations (ed R.A. Hinde) 61–74 (Cambridge University Press, 1969).

9 Price, P. H. Developmental determinants of structure in zebra finch song. Journal of Comparative and Physiological Psychology 93, 260–277 (1979).

10 Sossinka, R. & Bohner, J. Song types in the zebra finch Poephila guttata castanotis. Zeitschrift fur Tierpsychologie 53, 123–132 (1980).

11 Bohner, J. Song learning in the zebra finch (Taeniopygia guttata): selectivity in choice of a tutor and accuracy of song copies. Animal Behavior 31, 231–237 (1983).

12 Zann, R. A. Structural variation in the zebra finch distance call. Zeitschrift fur Tierpsychologie 66, 328–345 (1984).

13 Clayton, N. S. Song learning in cross-fostered zebra finches: A re-examination of the sensitive phase. Behaviour 102, 67–81 (1987).

14 Bohner, J. Early acquisition of song in the zebra finch, Taeniopygia guttata. Animal Behavior 39, 369–374 (1990).

15 Zann, R. A. The Zebra Finch: a Synthesis of Field and Laboratory Studies. (Oxford University Press, 1996).

16 Tchernichovski, O., Lints, T., Mitra, P. P. & Nottebohm, F. Vocal imitation in zebra finches is inversely related to model abundance. Proc Natl Acad Sci U S A 96, 12901–12904 (1999).

17 Tchernichovski, O., Mitra, P. P., Lints, T. & Nottebohm, F. Dynamics of the vocal imitation process: how a zebra finch learns its song. Science 291, 2564–2569 (2001).

18 Funabiki, Y. & Konishi, M. Long memory in song learning by zebra finches. J Neurosci 23, 6928–6935 (2003).

19 Goller, F. & Cooper, B. G. Peripheral motor dynamics of song production in the zebra finch. Ann N Y Acad Sci 1016, 130–152 (2004). 10.1196/annals.1298.009

20 Alam, D., Zia, F. & Roberts, T. F. The hidden fitness of the male zebra finch courtship song. Nature (2024). 10.1038/s41586-024-07207-4

21 Nottebohm, F., Stokes, T. M. & Leonard, C. M. Central control of song in the canary, Serinus canarius. Journal of Comparative Neurology 165, 457–486 (1976).

22 Konishi, M. Birdsong for neurobiologists. Neuron 3, 541–549 (1989).

23 Schmidt, M. F. & Goller, F. Breathtaking Songs: Coordinating the Neural Circuits for Breathing and Singing. Physiology (Bethesda) 31, 442–451 (2016). 10.1152/physiol.00004.2016

24 Schmidt, M. F., Ashmore, R. C. & Vu, E. T. Bilateral control and interhemispheric coordination in the avian song motor system. Ann N Y Acad Sci 1016, 171–186 (2004).

25 Mooney, R. Neural mechanisms for learned birdsong. Learn Mem 16, 655–669 (2009). 10.1101/lm.1065209

26 Fee, M. S., Kozhevnikov, A. A. & Hahnloser, R. H. Neural mechanisms of vocal sequence generation in the songbird. Ann N Y Acad Sci 1016, 153–170 (2004).

27 Roberts, T. F. et al. Identification of a motor-to-auditory pathway important for vocal learning. Nat Neurosci 20, 978–986 (2017). 10.1038/nn.4563

28 Aronov, D., Andalman, A. S. & Fee, M. S. A specialized forebrain circuit for vocal babbling in the juvenile songbird. Science 320, 630–634 (2008).

29 Moll, F. W. et al. Thalamus drives vocal onsets in the zebra finch courtship song. Nature 616, 132–136 (2023). 10.1038/s41586-023-05818-x

30 Daliparthi, V. K. et al. Transitioning between preparatory and precisely sequenced neuronal activity in production of a skilled behavior. eLife 8 (2019). 10.7554/eLife.43732

31 Stacho, M. et al. A cortex-like canonical circuit in the avian forebrain. Science 369 (2020). 10.1126/science.abc5534

32 Colquitt, B. M., Merullo, D. P., Konopka, G., Roberts, T. F. & Brainard, M. S. Cellular transcriptomics reveals evolutionary identities of songbird vocal circuits. Science 371 (2021). 10.1126/science.abd9704

33 Ikeda, M. Z., Trusel, M. & Roberts, T. F. Memory circuits for vocal imitation. Curr Opin Neurobiol 60, 37–46 (2020). 10.1016/j.conb.2019.11.002

34 Zhao, W., Garcia-Oscos, F., Dinh, D. & Roberts, T. F. Inception of memories that guide vocal learning in the songbird. Science 366, 83–89 (2019).

35 Bauer, E. E. et al. A synaptic basis for auditory-vocal integration in the songbird. J Neurosci 28, 1509–1522 (2008). 10.1523/jneurosci.3838-07.2008

36 Garcia-Oscos, F. et al. Autism-linked gene FoxP1 selectively regulates the cultural transmission of learned vocalizations. Science advances 7 (2021). 10.1126/sciadv.abd2827

37 Nottebohm, F., Kelley, D. B. & Paton, J. A. Connections of vocal control nuclei in the canary telencephalon. Journal of Comparative Neurology 207, 344–357 (1982).

38 Vates, G. E., Broome, B. M., Mello, C. V. & Nottebohm, F. Auditory pathways of caudal telencephalon and their relation to the song system of adult male zebra finches. Journal of Comparative Neurology 366, 613–642 (1996).

39 Akutagawa, E. & Konishi, M. New brain pathways found in the vocal control system of a songbird. Journal of Comparative Neurology 518, 3086–3100 (2010). 10.1002/cne.22383

40 Roberts, T. F., Klein, M. E., Kubke, M. F., Wild, J. M. & Mooney, R. Telencephalic neurons monosynaptically link brainstem and forebrain premotor networks necessary for song. J Neurosci 28, 3479–3489 (2008).

41 Foster, E. F. & Bottjer, S. W. Lesions of a telencephalic nucleus in male zebra finches: Influences on vocal behavior in juveniles and adults. J Neurobiol 46, 142–165 (2001). 10.1002/1097-4695(20010205)46:2<142::aid-neu60>3.0.co;2-r

42 Danish, H. H., Aronov, D. & Fee, M. S. Rhythmic syllable-related activity in a songbird motor thalamic nucleus necessary for learned vocalizations. PloS one 12, e0169568 (2017). 10.1371/journal.pone.0169568

43 Coleman, M. J. & Vu, E. T. Recovery of impaired songs following unilateral but not bilateral lesions of nucleus uvaeformis of adult zebra finches. J Neurobiol 63, 70–89 (2005).

44 Hamaguchi, K. & Mooney, R. Recurrent interactions between the input and output of a songbird cortico-basal ganglia pathway are implicated in vocal sequence variability. J Neurosci 32, 11671–11687 (2012). 10.1523/JNEUROSCI.1666-12.2012

45 Hamaguchi, K., Tanaka, M. & Mooney, R. A Distributed Recurrent Network Contributes to Temporally Precise Vocalizations. Neuron 91, 680–693 (2016). 10.1016/j.neuron.2016.06.019

46 Roberts, T. F., Gobes, S. M., Murugan, M., Olveczky, B. P. & Mooney, R. Motor circuits are required to encode a sensory model for imitative learning. Nat Neurosci 15, 1454–1459 (2012). 10.1038/nn.3206

47 Mardinly, A. R. et al. Precise multimodal optical control of neural ensemble activity. Nat Neurosci 21, 881–893 (2018). 10.1038/s41593-018-0139-8

48 Govorunova, E. G., Sineshchekov, O. A., Janz, R., Liu, X. & Spudich, J. L. NEUROSCIENCE. Natural light-gated anion channels: A family of microbial rhodopsins for advanced optogenetics. Science 349, 647–650 (2015). 10.1126/science.aaa7484

49 Malyshev, A. Y. et al. Chloride conducting light activated channel GtACR2 can produce both cessation of firing and generation of action potentials in cortical neurons in response to light. Neurosci Lett 640, 76–80 (2017). 10.1016/j.neulet.2017.01.026

50 Messier, J. E., Chen, H., Cai, Z. L. & Xue, M. Targeting light-gated chloride channels to neuronal somatodendritic domain reduces their excitatory effect in the axon. eLife 7 (2018). 10.7554/eLife.38506

51 Khirug, S. et al. GABAergic depolarization of the axon initial segment in cortical principal neurons is caused by the Na-K-2Cl cotransporter NKCC1. J Neurosci 28, 4635–4639 (2008). 10.1523/jneurosci.0908-08.2008

52 Linders, L. E. et al. Studying Synaptic Connectivity and Strength with Optogenetics and Patch-Clamp Electrophysiology. Int J Mol Sci 23 (2022). 10.3390/ijms231911612

53 Coleman, M. J., Roy, A., Wild, J. M. & Mooney, R. Thalamic gating of auditory responses in telencephalic song control nuclei. Journal of Neuroscience 27, 10024–10036 (2007). 10.1523/JNEUROSCI.2215-07.2007

54 Faunes, M. & Wild, J. M. The ascending projections of the nuclei of the descending trigeminal tract (nTTD) in the zebra finch (Taeniopygia guttata). J Comp Neurol 525, 2832–2846 (2017). 10.1002/cne.24247

55 Wild, J. M. & Gaede, A. H. Second tectofugal pathway in a songbird (Taeniopygia guttata) revisited: Tectal and lateral pontine projections to the posterior thalamus, thence to the intermediate nidopallium. J Comp Neurol 524, 963–985 (2016). 10.1002/cne.23886

56 Wild, J. M., Krutzfeldt, N. O. & Kubke, M. F. Connections of the auditory brainstem in a songbird, Taeniopygia guttata. III. Projections of the superior olive and lateral lemniscal nuclei. J Comp Neurol 518, 2149–2167 (2010). 10.1002/cne.22325

57 Wild, J. M. Visual and somatosensory inputs to the avian song system via nucleus uvaeformis (Uva) and a comparison with the projections of a similar thalamic nucleus in a nonsongbird, Columba livia. Journal of Comparative Neurology 349, 512–535 (1994).

58 Akutagawa, E. & Konishi, M. Connections of thalamic modulatory centers to the vocal control system of the zebra finch. Proc Natl Acad Sci U S A 102, 14086–14091 (2005). 10.1073/pnas.0506774102

59 Coleman, M. J. & Mooney, R. Synaptic transformations underlying highly selective auditory representations of learned birdsong. J Neurosci 24, 7251–7265 (2004).

60 Mackevicius, E. L., Happ, M. T. L. & Fee, M. S. An avian cortical circuit for chunking tutor song syllables into simple vocal-motor units. Nature communications 11, 5029 (2020). 10.1038/s41467-020-18732-x

61 Otchy, T. M. et al. Acute off-target effects of neural circuit manipulations. Nature 528, 358–363 (2015). 10.1038/nature16442

62 Fortune, E. S. & Margoliash, D. Parallel pathways and convergence onto HVc and adjacent neostriatum of adult zebra finches (Taeniopygia guttata). Journal of Comparative Neurology 360, 413–441 (1995).

63 Kelley, D. B. & Nottebohm, F. Projections of a telencephalic auditory nucleus-field L-in the canary. Journal of Comparative Neurology 183, 455–469 (1979).

64 Coleman M. J. M. R. Synaptic transformations underlying highly selective auditory representations of learned birdsong. J. Neurosci., Aug 18;24(33):7251–7265. doi: 7210.1523/JNEUROSCI.0947-7204.2004. (2004).

65 Koparkar, A. et al. (Cold Spring Harbor Laboratory, 2023).

66 Hickok, G. & Poeppel, D. The cortical organization of speech processing. Nat Rev Neurosci 8, 393–402 (2007). 10.1038/nrn2113

67 Poeppel, D., Emmorey, K., Hickok, G. & Pylkkanen, L. Towards a new neurobiology of language. J Neurosci 32, 14125–14131 (2012). 10.1523/JNEUROSCI.3244-12.2012

68 Hertrich, I., Dietrich, S. & Ackermann, H. The Margins of the Language Network in the Brain. Frontiers in Communication 5 (2020). 10.3389/fcomm.2020.519955

69 Simpson, H. B. & Vicario, D. S. Brain pathways for learned and unlearned vocalizations differ in zebra finches. J Neurosci 10, 1541–1556 (1990). 10.1523/jneurosci.10-05-01541.1990

70 Basista, M. J. et al. Independent premotor encoding of the sequence and structure of birdsong in avian cortex. J Neurosci 34, 16821–16834 (2014). 10.1523/JNEUROSCI.1940-14.2014

71 Vu, E. T., Mazurek, M. E. & Kuo, Y. C. Identification of a forebrain motor programming network for the learned song of zebra finches. J Neurosci 14, 6924–6934 (1994). 10.1523/jneurosci.14-11-06924.1994

72 Ashmore, R. C., Wild, J. M. & Schmidt, M. F. Brainstem and forebrain contributions to the generation of learned motor behaviors for song. J Neurosci 25, 8543–8554 (2005).

73 Vu, E. T., Schmidt, M. F. & Mazurek, M. E. Interhemispheric coordination of premotor neural activity during singing in adult zebra finches. J Neurosci 18, 9088–9098 (1998). 10.1523/jneurosci.18-21-09088.1998

74 Sanchez-Valpuesta, M. et al. Corticobasal ganglia projecting neurons are required for juvenile vocal learning but not for adult vocal plasticity in songbirds. Proc Natl Acad Sci U S A 116, 22833–22843 (2019). 10.1073/pnas.1913575116

75 Tanaka, M., Sun, F., Li, Y. & Mooney, R. A mesocortical dopamine circuit enables the cultural transmission of vocal behaviour. Nature 563, 117–120 (2018). 10.1038/s41586-018-0636-7

76 Koparkar, A. et al. Lesions in a songbird vocal circuit increase variability in song syntax. eLife 13 (2024). 10.7554/eLife.93272

77 Kosche, G., Vallentin, D. & Long, M. A. Interplay of inhibition and excitation shapes a premotor neural sequence. J Neurosci 35, 1217–1227 (2015). 10.1523/jneurosci.4346-14.2015

78 Vallentin, D., Kosche, G., Lipkind, D. & Long, M. A. Neural circuits. Inhibition protects acquired song segments during vocal learning in zebra finches. Science 351, 267–271 (2016). 10.1126/science.aad3023

79 Mooney, R. & Prather, J. F. The HVC microcircuit: the synaptic basis for interactions between song motor and vocal plasticity pathways. J Neurosci 25, 1952–1964 (2005).

80 Tremblay, R., Lee, S. & Rudy, B. GABAergic Interneurons in the Neocortex: From Cellular Properties to Circuits. Neuron 91, 260–292 (2016). 10.1016/j.neuron.2016.06.033

81 Berger, T. K., Silberberg, G., Perin, R. & Markram, H. Brief bursts self-inhibit and correlate the pyramidal network. PLoS Biol 8 (2010). 10.1371/journal.pbio.1000473

82 Kapfer, C., Glickfeld, L. L., Atallah, B. V. & Scanziani, M. Supralinear increase of recurrent inhibition during sparse activity in the somatosensory cortex. Nat Neurosci 10, 743–753 (2007). 10.1038/nn1909

83 Silberberg, G. & Markram, H. Disynaptic inhibition between neocortical pyramidal cells mediated by Martinotti cells. Neuron 53, 735–746 (2007). 10.1016/j.neuron.2007.02.012

84 Anastasiades, P. G., Marlin, J. J. & Carter, A. G. Cell-Type Specificity of Callosally Evoked Excitation and Feedforward Inhibition in the Prefrontal Cortex. Cell reports 22, 679–692 (2018). 10.1016/j.celrep.2017.12.073

85 Delevich, K., Tucciarone, J., Huang, Z. J. & Li, B. The mediodorsal thalamus drives feedforward inhibition in the anterior cingulate cortex via parvalbumin interneurons. J Neurosci 35, 5743–5753 (2015). 10.1523/JNEUROSCI.4565-14.2015

86 Bruno, R. M. & Sakmann, B. Cortex is driven by weak but synchronously active thalamocortical synapses. Science 312, 1622–1627 (2006). 10.1126/science.1124593

87 Bruno, R. M. & Simons, D. J. Feedforward mechanisms of excitatory and inhibitory cortical receptive fields. J Neurosci 22, 10966–10975 (2002). 10.1523/JNEUROSCI.22-24-10966.2002

88 Cardin, J. A., Kumbhani, R. D., Contreras, D. & Palmer, L. A. Cellular mechanisms of temporal sensitivity in visual cortex neurons. J Neurosci 30, 3652–3662 (2010). 10.1523/JNEUROSCI.5279-09.2010

89 Pinto, D. J., Brumberg, J. C. & Simons, D. J. Circuit dynamics and coding strategies in rodent somatosensory cortex. J Neurophysiol 83, 1158–1166 (2000). 10.1152/jn.2000.83.3.1158

90 Pinto, D. J., Hartings, J. A., Brumberg, J. C. & Simons, D. J. Cortical damping: analysis of thalamocortical response transformations in rodent barrel cortex. Cereb Cortex 13, 33–44 (2003). 10.1093/cercor/13.1.33

91 Dimidschstein, J. et al. A viral strategy for targeting and manipulating interneurons across vertebrate species. Nat Neurosci 19, 1743–1749 (2016). 10.1038/nn.4430

92 Stauffer, T. R. et al. Axial organization of a brain region that sequences a learned pattern of behavior. J Neurosci 32, 9312–9322 (2012). 10.1523/JNEUROSCI.0978-12.2012

93 Day, N. F., Terleski, K. L., Nykamp, D. Q. & Nick, T. A. Directed functional connectivity matures with motor learning in a cortical pattern generator. J Neurophysiol 109, 913–923 (2013). 10.1152/jn.00937.2012

94 Opitz, A., Falchier, A., Linn, G. S., Milham, M. P. & Schroeder, C. E. Limitations of ex vivo measurements for in vivo neuroscience. Proc Natl Acad Sci U S A 114, 5243–5246 (2017). 10.1073/pnas.1617024114

95 Ballanyi, K. & Ruangkittisakul, A. in Encyclopedia of Neuroscience (eds Marc D. Binder, Nobutaka Hirokawa, & Uwe Windhorst) 483–490 (Springer Berlin Heidelberg, 2009).

96 Ben-Tov, M., Duarte, F. & Mooney, R. A neural hub for holistic courtship displays. Curr Biol 33, 1640–1653 e1645 (2023). 10.1016/j.cub.2023.02.072

97 Xiao, L. et al. A Basal Ganglia Circuit Sufficient to Guide Birdsong Learning. Neuron 98, 208–221.e205 (2018). 10.1016/j.neuron.2018.02.020

